# Single Cell, Single Nucleus and Spatial RNA Sequencing of the Human Liver Identifies Hepatic Stellate Cell and Cholangiocyte Heterogeneity

**DOI:** 10.1101/2021.03.27.436882

**Authors:** Tallulah S. Andrews, Jawairia Atif, Jeff C. Liu, Catia T. Perciani, Xue-Zhong Ma, Cornelia Thoeni, Michal Slyper, Gökcen Eraslan, Asa Segerstolpe, Justin Manuel, Sai Chung, Erin Winter, Iulia Cirlan, Nicholas Khuu, Sandra Fischer, Orit Rozenblatt-Rosen, Aviv Regev, Ian D. McGilvray, Gary D. Bader, Sonya A. MacParland

## Abstract

The critical functions of the human liver are coordinated through the interactions of hepatic parenchymal and non-parenchymal cells. Recent advances in single cell transcriptional approaches have enabled an examination of the human liver with unprecedented resolution. However, dissociation related cell perturbation can limit the ability to fully capture the human liver’s parenchymal cell fraction, which limits the ability to comprehensively profile this organ. Here, we report the transcriptional landscape of 73,295 cells from the human liver using matched single-cell RNA sequencing (scRNA-seq) and single-nucleus RNA sequencing (snRNA-seq). The addition of snRNA-seq enabled the characterization of interzonal hepatocytes at single-cell resolution, revealed the presence of rare subtypes of hepatic stellate cells previously only seen in disease, and detection of cholangiocyte progenitors that had only been observed during *in vitro* differentiation experiments. However, T and B lymphocytes and NK cells were only distinguishable using scRNA-seq, highlighting the importance of applying both technologies to obtain a complete map of tissue-resident cell-types. We validated the distinct spatial distribution of the hepatocyte, cholangiocyte and stellate cell populations by an independent spatial transcriptomics dataset and immunohistochemistry. Our study provides a systematic comparison of the transcriptomes captured by scRNA-seq and snRNA-seq and delivers a high-resolution map of the parenchymal cell populations in the healthy human liver.

## INTRODUCTION

The liver is an essential organ responsible for critical functions including lipid and glucose metabolism, protein synthesis, bile secretion and immune functions. Single cell RNA sequencing (scRNA-seq) technologies enable the analysis of the transcriptome of individual cells and have provided important insights regarding development(1), physiology(2,3) and pathology(4–6) of the human liver. These studies have shed light into previously inaccessible aspects of human liver physiology such as hepatic lobular zonation, cell to cell interactions, and immune cell phenotype and heterogeneity.

Previously, we examined the cellular complexity of the human liver by single cell RNA sequencing (scRNA-seq) and identified 20 distinct cell clusters including two distinct populations of liver resident macrophages with immunoregulatory and inflammatory properties.(2) An observation from this work was that enzymatic and mechanical dissociation of the human liver tissue significantly impacted the composition of the liver map in that hepatocytes were sensitive to dissociation-induced damage and cholangiocytes and hepatic stellate cells were not well-released by our dissociation technique. For example, cholangiocytes, parenchymal cells that form the bile duct and are expected to make up 3-5% of all liver cells,(7) comprised only 199 (0.64%) cells of our scRNA-seq 8444 map. Capturing cholangiocyte heterogeneity is key to understanding the pathogenesis of cholangiopathies, such as primary sclerosing cholangitis, for which there are no curative therapeutic interventions.(8)

Single nucleus RNA sequencing (snRNA-seq) is an approach that bypasses the cell dissociation step required for scRNA-seq by using detergents to release nuclei from intact cells. SnRNA-seq data is also compatible with snap frozen samples that may be available from tissue archives. Recently, Slyper et al.(9) assessed three different nuclei isolation protocols for snRNA-seq from frozen tissues that each employed different detergent based buffers: Nonidet P40 with salts and Tris (NST), Tween-20 with salts and Tris (TST), and CHAPS with salts and Tris (CST). Here, we carried out matched snRNA-seq using these three protocols and scRNA-seq using our published experimental and analysis workflow(2) on four healthy human liver samples (Fig. 1a). Using multiple protocols on the same samples enables us to evaluate the ability of these three snRNA-seq protocols to reduce dissociation-related effects compared to scRNA-seq protocols, contrast the expression profiles of cells measured with each protocol, and develop an approach to integrate the results into a single map that is more comprehensive than possible with any individual method.

**Fig. 1:**
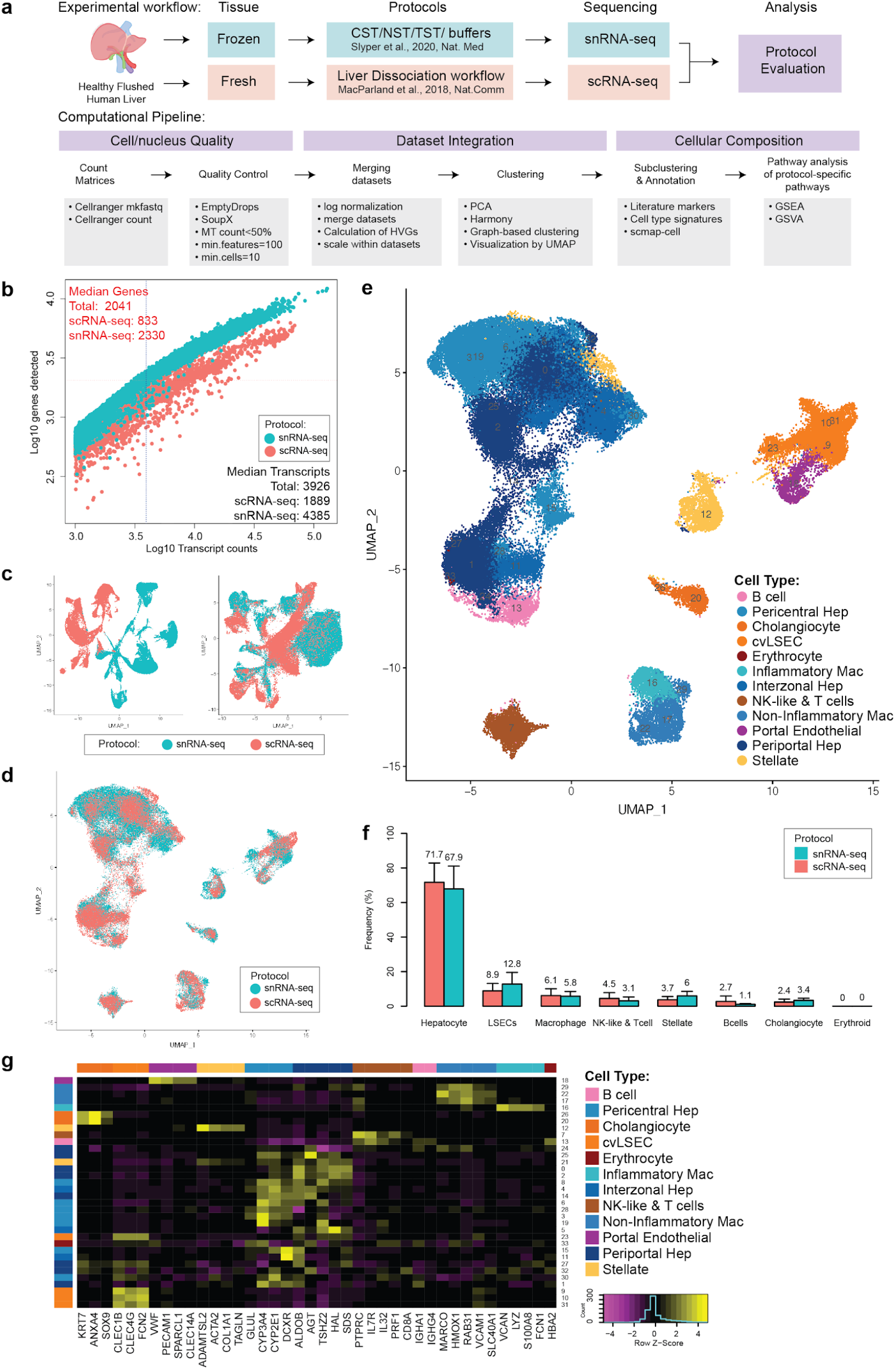
Technical differences between scRNA-seq and snRNA-seq in profiling cells from the healthy human liver. **a:** Overview of single cell and single nucleus isolation, datasets integration and analysis workflows. **b:** Sensitivity of each approach as measured by the number of genes and transcripts identified in each cell/nucleus. **c:** UMAP projection of cells derived from scRNA-seq and snRNA-seq i) merged and then scaled ii) individually scaled before merging. **d:** UMAP projection of cells from scRNA-seq (pink) and snRNA-seq (blue) individually scaled before merging and then integrated using harmony. **e:** UMAP plot showing the assigned identity for each cluster after scaling individually, merging and integrating. **f:** Frequency of each major cell population in their source dataset, error bars indicates 95% confidence intervals across samples. **g:** Heatmap showing scaled mean expression of known marker genes in each cluster. LSECs: Liver sinusoidal endothelial cells, cvLSECS: Central Venous Liver Sinusoidal endothelial cells. UMAP: Uniform Manifold Approximation and Projection. PCA: Principal Component Analysis

Our work highlights cell type composition differences between snRNA-seq and scRNA-seq technologies as applied to human liver, and reveals cholangiocyte and hepatic stellate cell subpopulations specific to the snRNA-seq data, previously not identified in single cell transcriptomic studies. Combining results from both technologies creates a rich new data set for the interpretation of human liver biology, identifying key cell-type defining marker genes across both technologies (Supplementary Table 1).

## RESULTS

### Examining the quality of liver mapping *via* scRNA-seq *vs*. scRNA-seq

Single cell and single nucleus transcriptomes were generated from four healthy human livers from neurologically deceased donors that were undergoing transplantation into living recipients. Samples were chosen from a total of 23 to preference those with the most intact hepatocytes in preliminary analyses. In total, 29,432 single cells were captured from fresh liver dissociates and 43,863 single nuclei were captured from matched snap frozen tissue and sequenced using the 10X Chromium platform. These data underwent identical quality control processing, and the matched samples were systematically compared.

SnRNA-seq captured a greater diversity of genes than scRNA-seq (Fig. 1b). These differences are largely due to the high proportion of UMIs in scRNA-seq data that are derived from transcripts encoding ribosomal proteins and genes encoded in the mitochondrial genome which are not present in snRNA-seq data (Supplementary Table 2). Minimal differences were observed between different detergents used to extract nuclei (Supplementary Table 3). Furthermore, snRNA-seq contained a significantly higher proportion of ambient RNA than scRNA for most samples, estimated by SoupX (Supplementary Fig. 1).

### Integrating scRNA-seq and snRNA-seq maps

Using a typical computational processing pipeline, scRNA and snRNA do not cluster together (Fig. 1c, Extended Data Fig. 1), due to the systematic differences between RNA found in the nucleus *versus* the cytoplasm of cells. Additional technical confounding effects may be introduced during tissue processing and cell handling. The scRNA-seq samples were derived from fresh tissue that was enzymatically and mechanically dissociated which may introduce stress responses in cells. Whereas, the snRNA-seq samples were extracted from flash frozen tissue which should be less impacted by dissociation-related stresses. In addition, we see significant batch effects between individual donors when using the same sequencing technology, particularly in hepatocytes (Extended Data Fig. 2). This may be related to environmental influences on liver metabolism, and is consistent with our previously reported single-cell liver map.(2)

However, if samples are normalized and scaled individually before merging, cells and nuclei broadly cluster by cell-type rather than by transcriptome mapping technology (Fig. 1c). However, significant differences between technologies are still evident and application of Harmony,(10) a commonly used single cell data integration method, overcomes this and enables integrationed and co-clustering of scRNA-seq and snRNA-seq data (Fig. 1d).

### Systematic differences in nuclear and whole cell transcriptomes

The necessity to scale datasets individually to enable scRNA-seq and snRNA-seq to be integrated demonstrates that there are significant systematic gene expression differences between data generated by these technologies. Examining these systematically differentially expressed genes revealed that both gene function and gene length were strongly associated with expression as measured by scRNA-seq or snRNA-seq (Extended Data Fig. 3, 4). Genes encoded by the mitochondrial genome and nuclear genes that encode mitochondrial proteins are more highly expressed in scRNA-seq (Extended Data Fig. 4). Similarly, mRNA encoding ribosome-related proteins are more than 4-fold more highly expressed in scRNA-seq than snRNA-seq. This is expected, as mitochondria and ribosomes are prevalent in the cytoplasm in liver cells, especially hepatocytes, and cytoplasm is mostly missing from snRNA-seq material input. In contrast, other “housekeeping” protein-coding genes(11) were equally expressed in snRNA-seq and scRNA-seq. In agreement with results from other tissues,(12,13) long non-coding RNAs (lncRNA) were 0.7-fold more highly expressed in snRNA-seq than scRNA-seq, however, this is not significantly different from other non-housekeeping protein-coding genes, which were 0.8-fold more highly expressed in snRNA-seq.

Aside from gene function, overall gene length had the strongest correlation with relative expression in snRNA-seq *vs*. scRNA-seq (Extended Data Fig. 5). Longer genes (>37kb) were more highly expressed in snRNA-seq, and short genes (<15kb) were more highly expressed in scRNA-seq. This was true even after excluding ribosomal and mitochondrial-related genes.

### The cells of the human liver as revealed by scRNA-seq vs. snRNA-seq

After integrating the data (see: Methods), the data was clustered and annotated using known markers;(2) the resulting map revealed all major known hepatic cell-types (Fig. 1e,g). These cell types were represented in both scRNA-seq and snRNA-seq and in all samples (Extended Data Fig. 5,6), but were captured at different frequencies. In particular scRNA-seq captured a higher proportion of immune cells with 7% of all cells sequenced identified as lymphocytes and 9% identified as macrophages, compared to snRNA-seq with 3% lymphocytes and 5% macrophages. In contrast, snRNA-seq captured twice as many cholangiocytes, and hepatic stellate cells as scRNA-seq (Fig. 1f, Supplementary Table 4). We obtained similar percentages of liver sinusoidal endothelial cells (LSECs) and endothelial cells with both methods. However, these frequencies depend on the detergent used for the snRNA-seq. CST and NST extracted a higher frequency of LSECs, 25% and 20% respectively, and higher frequencies of stellate cells, 7% in CST and 12% in NST, but lower frequencies of hepatocytes.

### Hepatocytes

Hepatocytes are the main parenchymal cell of the hepatic lobule responsible for the majority of liver function. They exhibit functional zonation from the pericentral vein to the periportal region. Currently it is difficult to identify interzonal human hepatocytes due to a paucity of known markers.(2,3) Recent work has demonstrated the importance of hepatocytes of this region for liver homeostasis and regeneration.(14) Subclustering our hepatocyte cluster revealed six distinct clusters (Fig. 2a,b) sourced from both scRNA-seq and snRNA-seq (Fig. 2c), and all samples (Fig. 2d). Correlating these clusters with zonated expression from microdissected mice(15) enabled the annotation of three of these clusters as containing either pericentral (CV) or periportal hepatocytes (PP1, PP2) (Fig 2e). These annotations were confirmed using known pericentral marker genes (*CYP3A4, ADH4, GLUL*, and *BCHE*) and periportal marker genes (*HAL, CPS1*, and *HMGCS1)* (Fig. 1g). Of the remaining three subclusters, one was most strongly correlated with mouse layer 7, another was most strongly correlated with mouse layer 5 suggesting these two clusters (IZ1, IZ2) represent human interzonal hepatocytes (Fig. 2e). We validated the interzonal hepatocyte identity by comparing to bulk RNAseq derived from human microdissected livers, where these clusters correlated most strongly with interzonal layer 4(Extended Data Fig. 7).(5) Novel interzonal markers identified from these two clusters (*HINT1, COX7C, APOC1, FABP1, MT2A, MT1G* and *NDUFB1*) were validated using spatial transcriptomics and immunofluorescence where they exhibit clearly distinct expression patterns from either periportal or pericentral markers (Fig. 2g-i, Extended Data Fig. 8-12).

**Fig. 2:**
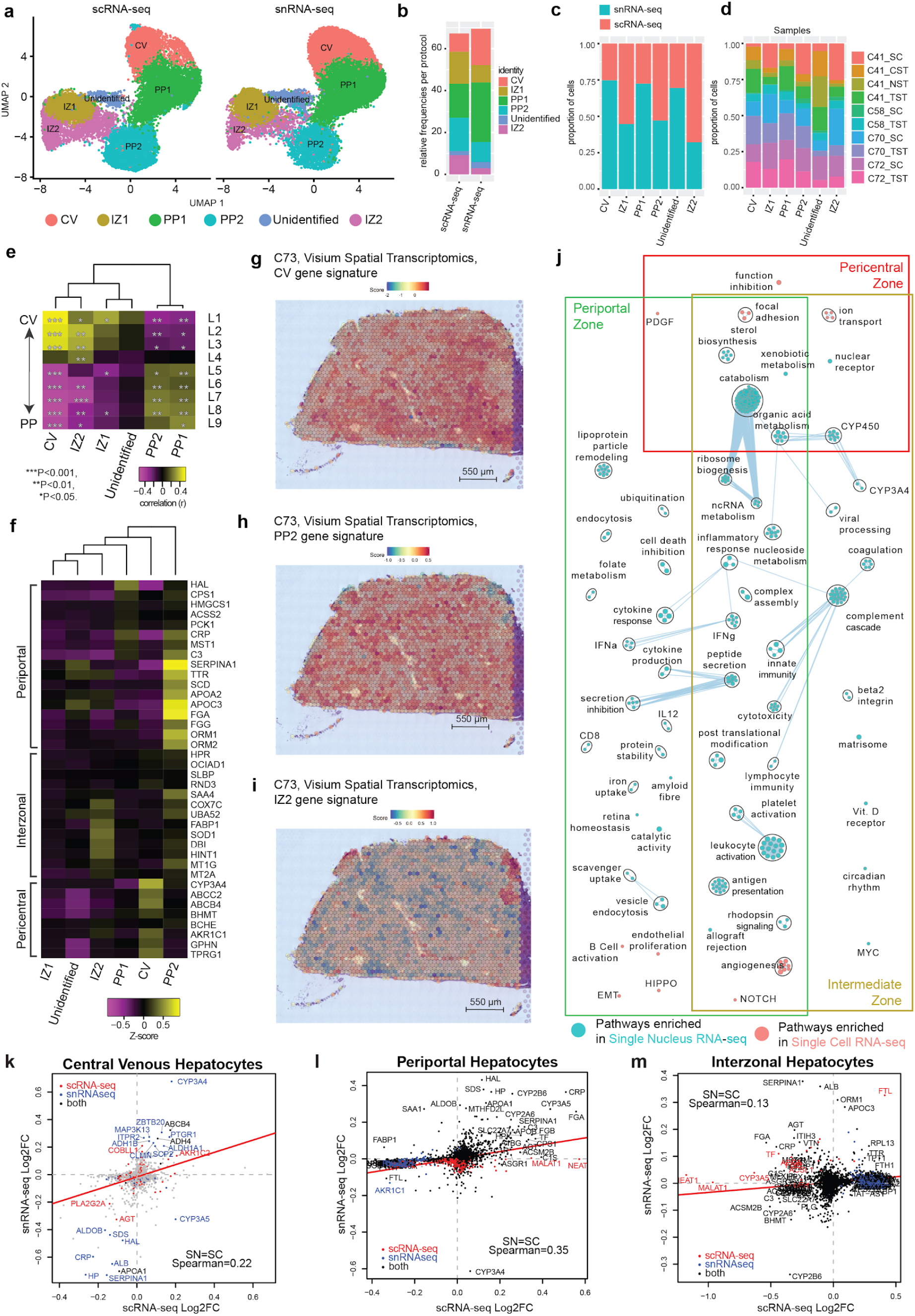
Hepatocyte populations in sample-matched scRNA-seq and snRNA-seq data are spatially resolved by spatial transcriptomics. **a:** UMAP plot with the six major populations of hepatocytes split by protocol. **b:** stacked bar-plot indicating the frequency of each population in either scRNA-seq or snRNA-seq datasets. Distribution of hepatocytes by protocol (**c**) or by donor sample (**d**) in the combined dataset. **e:** Correlation of human hepatocyte clusters to known mouse liver sinusoid layers calculated using Spearman correlation. \*\*\**P*<0.001, \*\**P*< 0.01, \**P*<0.05. **f:** Expression of known marker genes in hepatocyte subpopulations in the combined dataset. Gene signature scores of the top 30 marker genes in clusters CV (**g**), PP2 (**h**) and IZ2 (**i**) across the spatial transcriptomics spots of a healthy human liver cryosection. **j:** Pathway enrichment analysis examining which cellular pathways are better represented by snRNA-seq (cyan) and scRNA-seq (pink) in the central venous, periportal and interzonal hepatocyte populations. Circles (nodes) represent pathways, sized by the number of genes included in that pathway. Related pathways, indicated by light blue lines, are grouped into a theme (black circle) and labeled. Intra-and inter-pathway relationships are shown in light blue and represent the number of genes shared between each pathway. Log2FC of significant genes (q-value<0.05) within either scRNA-seq (Red) or snRNA-seq (blue) or both (black) for CV hepatocytes (**k: cluster CV**), PP clusters (**l:** clusters: PP1 and PP2), and IZ clusters (**m:** clusters: IZ1 and IZ2).CV: Central Venous, IZ: Inter-zonal, PP: Periportal.

The final cluster of *ALB-*expressing hepatocytes expressed periportal hepatocyte markers like *SERPINA1, TTR, APOA1*, and *APOC3*. However, this cluster did not correlate strongly with the periportal mouse sinusoid regions nor express many other periportal marker genes like *HAL, CPS1*, and *HMGCS1*. Furthermore, the expression profile correlated most strongly with the interzonal layer 4 of the human sinusoid, but did not correlate with any mouse zonation layer (top DE: *ALB, SERPINA1, APOA1, HP, FTL, APOC3, SAA1, STAB1, TTR, B2M, LIFR)*. These results suggest this cluster may represent a human-specific interzonal hepatocyte cluster, but further work will be required to confirm this identity.

Hepatocytes as a whole were captured equally well in sc and snRNA-seq (Extended Table 2), however CV1 hepatocytes were almost twice as frequent (17% vs 9%, p < 10^−30^) in snRNA-seq than scRNAseq (Fig. 2b,c, Extended Table 3). Whereas, interzonal hepatocytes were most frequent in scRNAseq (15% and 9% vs 8% and 3%, p < 10^−15^). Several significant genes identified in CV1 hepatocytes by snRNA-seq were not identified by scRNA-seq (Fig. 2k) while the gene expression patterns for periportal hepatocytes correlated significantly across both technologies (Fig. 2m). The greatest discordance exists for the interzonal hepatocytes where many of the genes upregulated in snRNA-seq are downregulated in scRNA-seq (Fig. 2l).

However, despite these differences we observe almost all hepatocyte related pathways exhibiting elevated expression in snRNA-seq-derived hepatocytes (Fig. 2j). This may reflect poor viability or disrupted cellular state within hepatocytes due to the dissociation(16) required for scRNAseq which is not present when examining hepatocytes with snRNA-seq. Thus, snRNA-seq may provide better characterization of hepatocytes than scRNAseq despite similar capture rates.

### Cholangiocytes

Cholangiocytes are epithelial cells that line the bile ducts and generate 30% of the total bile volume.(17) Our previous attempt to characterize these cells using exclusively scRNA-seq identified only a single population encompassing 1.4% (199/8444) of the cells expressing cholangiocyte markers (*EPCAM, SOX9* and *KRT1*).

In this study, we found that snRNA-seq captured a higher proportion of *KRT7, SOX9, ANXA4* expressing cholangiocytes (3.4% vs 2.4%) resulting in a total of 448 cholangiocyte-like cells (Fig. 1g, Extended Table 2). Subclustering this population revealed six transcriptionally-distinct subpopulations (labelled Chol-1 to Chol-6) (Fig. 3a). We identified three *ASGR1*+ hepatocyte-like clusters, two typical cholangiocyte-like clusters (*KRT7, KRT18, SLC4A2* high), and a cluster of progenitors,>82% of which were derived from snRNA-seq (Fig. 3b,c, Supplementary Table 6). These clusters were not specific to a particular sample or donor indicating they were not the result of batch effects or donor-specific variation (Fig. 3d). Rather, these clusters formed a branching trajectory extending from bipotent progenitors to both hepatocyte and cholangiocyte cell fates, as computed using both Slingshot(18) and diffusion maps (Fig. 3g, Extended Fig. 20).

**Fig. 3:**
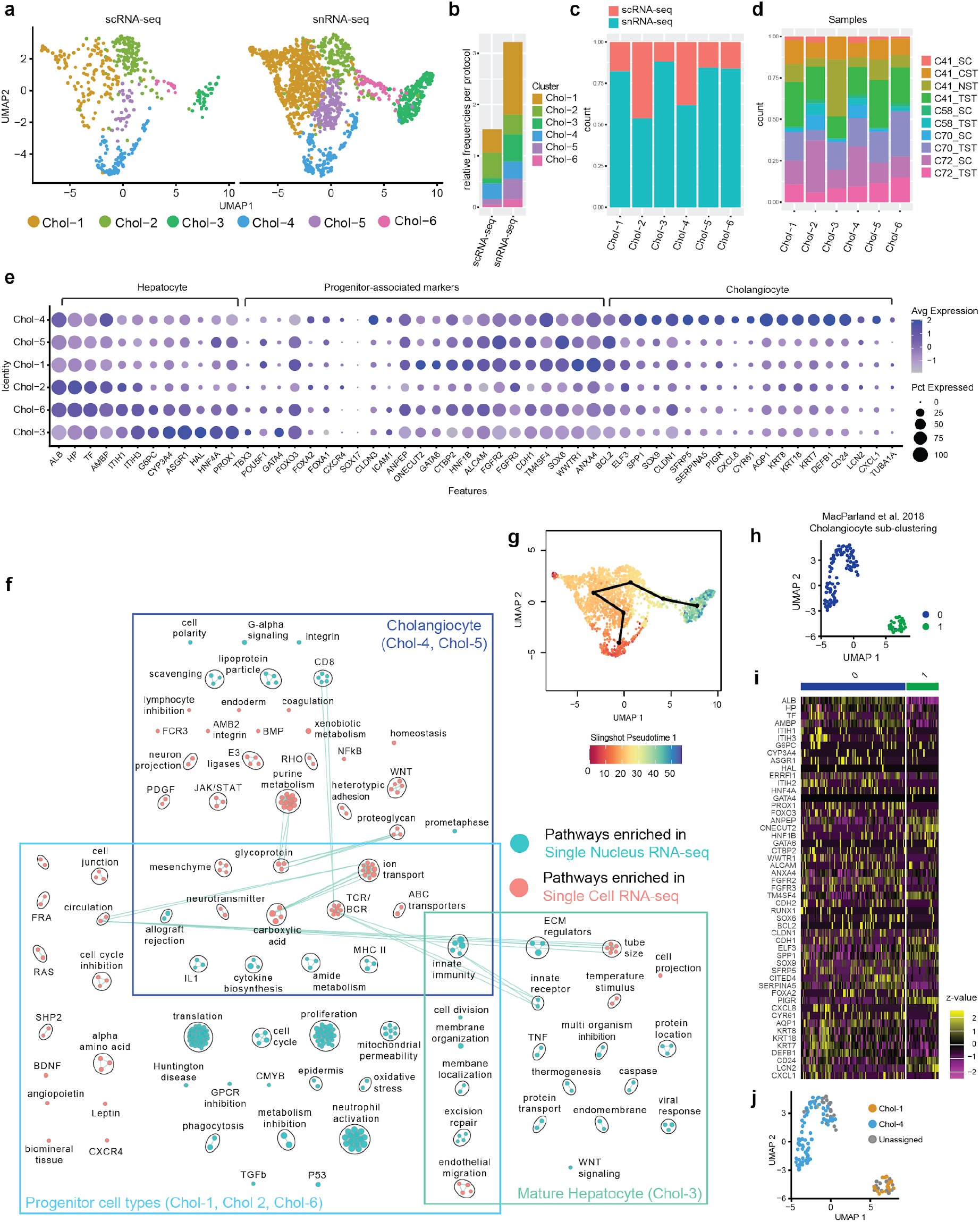
Cholangiocyte-associated cells as revealed by SnRNA-seq. **a:** UMAP plot with the six major populations of cholangiocytes and progenitor cells identified in the combined dataset split by protocol. **b:** Frequency of each population in either scRNA-seq or snRNA-seq datasets. Distribution of each population by protocol (**c**) and by sample (**d**) in the combined dataset. **e:** Expression of known cholangiocyte, progenitor and hepatocyte marker genes in each population. The size of the circle indicates the percentage of cells in each population expressing each gene. **f:** Pairwise pathway analysis comparing cholangiocyte-like, progenitor-like and mature hepatocyte-like cells from snRNA-seq (cyan) to those from scRNA-seq (pink). Circles (nodes) represent pathways, sized by the number of genes included in that pathway. Related pathways, indicated by light blue lines, are grouped into a theme (black circle) and labeled. Due to low cell number, similar clusters were combined for pathway analysis (**see Extended Data Fig. 15**) for more details. **g:** UMAP plot coloured by Slingshot (18) pseudotime values with the inferred cholangiocyte to hepatocyte trajectory overlayed. **h**: UMAP of sub-clustered cholangiocyte-like populations from an independent scRNA-seq healthy human liver dataset (2). **i:** Heatmap depicting the relative expression of cholangiocyte sub-population associated marker genes in (2) cholangiocyte-like cells. **j**: Projection of cell-type annotations from the combined scRNA-seq and snRNA-seq cholangiocyte dataset on to scRNA-seq data from (2) using scmap-cell (23). CV: Central Venous.

At the most differentiated end of the cholangiocyte branch was the Chol-4 population which contained mature cholangiocytes expressing differentiated cholangiocyte markers (*AQP1, KRT8, KRT18, KRT7, DEFB1, CD24, PIGR, ANXA4)* (Fig. 3e, Extended Data Fig. 13a), many of which are highly specific to human bile ducts (Extended Data Fig. 14a).(19) Furthermore, classical cholangiocyte pathways, such as cell polarity, ion transport, ABC transporters and many immune pathways, were enriched among the genes upregulated in this cluster (Fig. 3f, Extended Data Fig. 15a). Cells of this cluster were derived in equal measure from scRNA-seq and snRNA-seq, and the significantly upregulated markers, including *AQP1, SPP1* and *DEFB1*, were relatively consistent across both technologies (Extended Data Fig. 16). Thus mature cholangiocytes are well characterized by either sn-or scRNA-seq.

The less differentiated cholangiocyte population (Chol-5) was specific to snRNA-seq (177 nuclei *vs* 33 cells). Cholangiocyte identity was confirmed by the expression of key transcription factors (*HNF1B, ONECUT1*, and *SOX9)* and enrichment of WNT signalling, ABC and ion transporters(20)(Extended Data Fig. 15b). We determined this cluster contained specifically small cholangiocytes on the basis of *BCL2* expression (Fig. 3e, Extended Data Fig. 13a), which is not expressed by large cholangiocytes.(20) This was confirmed by noting bile-duct restricted expression of *BCL2* by immunohistochemistry(19) (Extended Data Fig. 14b). We note high expression of many stem-ness markers(21) in this cluster which is consistent with previous reports of a less differentiated phenotype of these cholangiocytes(20)(Fig. 3e, Extended Data Fig. 13a,b). Novel markers of this cluster were identified using snRNA-seq (*CYP3A5, PPARGC1A, FHIT, FMO5, TDO2, SOX6*), however, these were not recapitulated in scRNA-seq reinforcing that this population can only be characterized using snRNA-seq (Extended Data Fig. 16).

At the far end of the opposite branch was Chol-3, a population of central venous hepatocyte-like cells, expressing high levels of *CYP3A4, GPC6*, and *AOX1* (Extended Data Fig. 9). These cells clustered together with cholangiocytes because of their high expression of many bile metabolism genes (i.e. *ABCA8, ABCA6, HAO1, ABCA1, SLC27A2)* indicating involvement in biliary function. Interestingly, these cells express *HNF4a*, which is first expressed in progenitor hepatocytes and is a central regulator of hepatocyte differentiation.(22)

Two additional hepatocyte-biased clusters, Chol-2 and Chol-6, expressed some typical cholangiocyte markers (*KRT7, SOX9* and *CD24*) but also expressed early hepatocyte lineage-defining transcription factors (*HNF4a, FOXA2, PROX1, ONECUT1*, and *ONECUT2*) suggesting an intermediate or progenitor phenotype (Fig. 3e, Extended Data Fig. 13b,c, 17). Both Chol-6 and Chol-2 highly expressed genes related to progenitor-associated markers, such as *FOXA2, FGFR3, HES1, JAG1* (Fig. 3e, Extended Data Fig. 13b,d). The key difference between the clusters being the expression of proliferation-related genes in Chol-6. Indicating these cells were proliferative and non-proliferative hepatic progenitors.

At the root of these lineage-biased branches was a large cluster of bipotent progenitors (Chol-1), containing predominantly sequenced single-nuclei (621 nuclei *vs* 134 cells, Extended Table 6, Fig.3c). This cluster did not express mature cholangiocyte markers, rather a collection of many stem-like and progenitor markers, including *POU5F1, FOXO2, RUNX2, SOX6, CD133, ANPEP*, and *SOX9* (Extended Data Fig. 13b-d). This cluster was consistent with bipotent progenitors that have previously been observed only *in vitro*.(3) Furthermore, using pathway analysis we identify the NOTCH2 signalling pathway (*NOTCH2, RBPJ, MAML2, MAML3, MAML4)* as a key feature of these progenitor cells. Endoderm, cell cycle, and cell division pathways were enriched in this population relative to the other clusters supporting a stem-like state (Extended Data Fig. 14f, 17). Marker genes of Chol-1 (ie. *FOXO3, GATA6, FGFR3, ANPEP*) were localized to bile ducts in both our spatial transcriptomics data and publicly available immunofluorescence data, suggesting these may be the progenitor niche (Extended Data Fig. 18, 19).(19)

Employing the combined map presented here as a reference, we were able to use the automatic annotation tool scmap-cell(23) to identify and label two distinct subsets of cholangiocytes that were unable to be detected in our previous map (Fig 3i,j). We identified cells from both mature cholangiocytes (Chol-4) and a small number of bipotent progenitors (Chol-1) which could only be identified using the snRNA-seq data, demonstrating the utility of our combined sc & sn liver map.

### Hepatic stellate cells

Hepatic stellate cells (HSCs) are heterogeneous cells that can exist in either an activated or quiescent state and play a role in retinol storage and the response to liver injury.(24) Under physiological conditions, HSCs maintain a quiescent and non-proliferative phenotype and are localized between the layers of LSECs and hepatocytes in the space of Disse.(2,24). Previous single-cell studies have focused on heterogeneity in HSCs during disease or injury, and typically must use additional cell handling steps to enrich for HSCs.(4,25) However, using snRNA-seq we reveal that stellate cell heterogeneity is present in healthy livers at low frequencies (Fig 4). Subclustering our HSC cluster revealed seven distinct subtype of HSCs (HSC 1-7) present in healthy liver, of these only HSC-1, 2, and 4 contained more than 10 cells captured with scRNAseq and only HSC-4 was roughly equally captured by both methodologies and in all samples (Fig. 4a,c,d Extended Table 7). Overall, HSCs were composed of 2.5% of single nuclei, more than double their 1% of single cells captured (Fig. 4b).

**Fig. 4:**
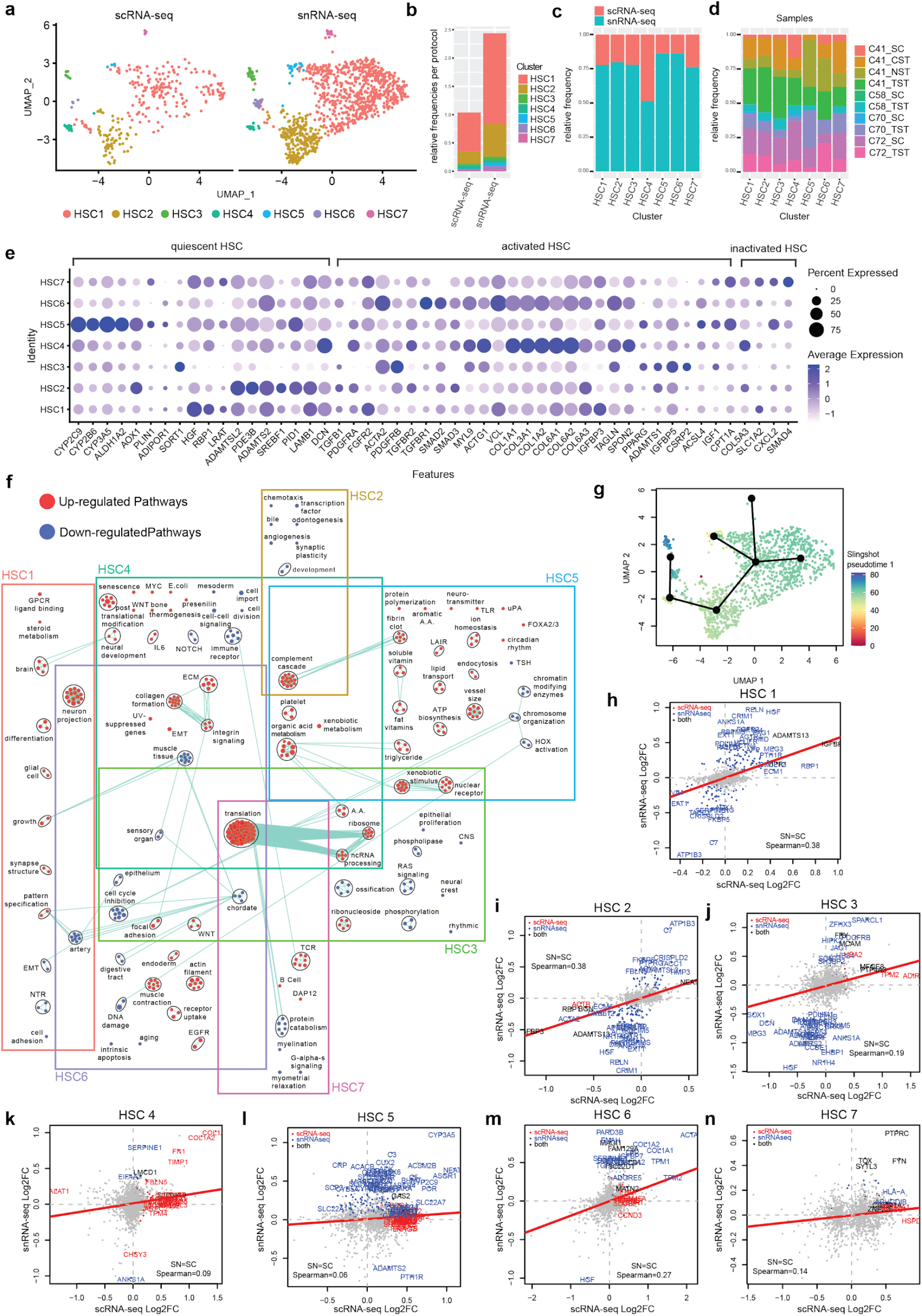
Identification of quiescent, activated and inactivated HSCs in the healthy human liver through scRNA-seq and snRNA-seq. **a:** UMAP plots with the seven major clusters of stellate cells in the combined dataset, split by protocol. **b:** stacked bar-plot indicating the frequency of each population in either scRNA-seq or snRNA-seq datasets. Distribution of each population by protocol (**c**) and by sample (**d**) in the combined dataset. **e:** Dot-plot indicating the relative expression of known stellate subtype specific marker genes in each population. The size of the circle indicates the percentage of cells in each cluster expressing each gene. **f**: Pathway enrichment analysis examining what are the upregulated (red) and down-regulated (cyan) biological pathways in each stellate cell cluster in the combined scRNA-seq and snRNA-seq dataset. Circles (nodes) represent pathways, sized by the number of genes included in that pathway. Related pathways, indicated by light blue lines, are grouped into a theme (black circle) and labeled. **g:** The original stellate cell UMAP plot coloured by Slingshot (18) pseudotime values with the inferred lineage overlayed. **h-n:** Log2FC of significant genes (q-value<0.05) within either scRNA-seq (Red) or snRNA-seq (blue) or both (black) for each cluster within the stellate cell dataset. HSC: Hepatic Stellate Cell.

The majority of HSCs (70% of HSC nuclei) comprised quiescent HSCs (HSC-1) specializing in vitamin A storage and metabolism with high expression of *RBP1, LRAT* and *PDE3B* (Extended Data Fig. 21a).(26) HSC initiation is characterized by the loss of retinol-bearing lipid droplets, the rapid induction of growth factor receptors, the development of a contractile phenotype and the modulation of growth factor signaling.(27) The second most frequent population (24% of HSC nuclei) were initiated HSCs (HSC-2) expressing retinol and lipid processing genes (*AOX1, PDE3D*, and *PDE4D*), as well as engaging in the complement cascade but lacking typical quiescence and activation markers (Fig. 4f). Furthermore, this cluster also showed expression of cytokine and growth factor receptors, and fibrotic, angiogenic and proliferative factors (i.e. *TGFB1, TGFBR2*) (Extended Data Fig. 21c,d).

The other 5 HSC clusters comprised only 10% of HSC nuclei (112/1069), and contained heterogeneous subsets of activated HSCs. HSCs typically become activated following liver damage, in response to cytokines, local damage, growth factors and fibrogenic signals.(24,28) HSC-4 and HSC-6 exhibited a pro-fibrogenic phenotype, expressing several fiber-associated genes: *ACTA2, SPARC, TAGLN, COL1A1, TIMP1* (Fig. 4e, Extended Data Fig. 21b). Enriched pathways included: collagen formation, extracellular matrix and integrin signaling in both of these clusters (Fig. 4f). HSC-6 expressed high levels of fibrogenic growth factors like *CTGF* and *TGFB1* (Extended Data Fig. 21d) and enrichment in contractile phenotype indicating a specialization in contractile fibers. Whereas HSC-4 was enriched in senescence-related pathways and inflammatory genes (*IL32, CSF1, TNFSF10, CCL2, IL6ST*) indicating a later stage in HSC activation. HSC-3 and HSC-5 expressed aHSC-associated genes *ACTA2* and *TAGLN* but not collagen or matrix remodelers. Importantly these clusters expressed high levels of *PPARG*, a transcription factor linked with reversion of activated HSC to a more quiescent phenotype, indicating these clusters contain iHSCs in the process of returning to a qHSC state (Extended Data Fig. 21e).(24,29) *RBP1, HGF, LRAT* and *DCN* gene expression and many lymphocyte-associated markers indicating they are likely to be doublets.

The trajectories inferred by both Slingshot and diffusion maps was consistent with these labels, estimating a path from qHSCs (HSC5, HSC1), through the aHSCs (HSC4 and HSC6), and ending at the iHSCs (HSC3) (Fig. 4g, Extended Data Fig. 21f,g). In addition, spatial transcriptomics independently confirmed qHSC expression to be higher and dispersed throughout liver tissue whereas aHSCs were primarily located in periportal regions (Extended Data Fig. 22). Only the qHSC population would be discoverable using scRNAseq alone, it is only because of the addition of snRNA-seq that the full diversity of HSCs in normal tissue was captured.

### Liver endothelial cells

The endothelium of the liver vasculature is made up of LSECs and vascular endothelial cells. LSEC populations were annotated using previously defined markers (Fig. 5a,e).(2) Similar to hepatocytes, central venous LSECs were more frequently captured with snRNA-seq (7.52%) than scRNA-seq (5.2%) (Fig. 5b-d). Whereas periportal LSECs and Portal endothelial cells were present in similar frequencies using either technology (ppLSECs: 1.15% in snRNA-seq and 1.32% in scRNA-seq, PortalEndo: 0.76% and 0.63% respectively, Supplementary Table 8). Markers of these populations were generally consistent across sc & snRNA-seq though fold-changes were typically smaller in snRNA-seq (Fig. 5g-i), this was particularly true for portal markers: *VWF, CLEC14A, ID1, SPARCL1, CTGF*. (Fig. 5e).

**Fig. 5:**
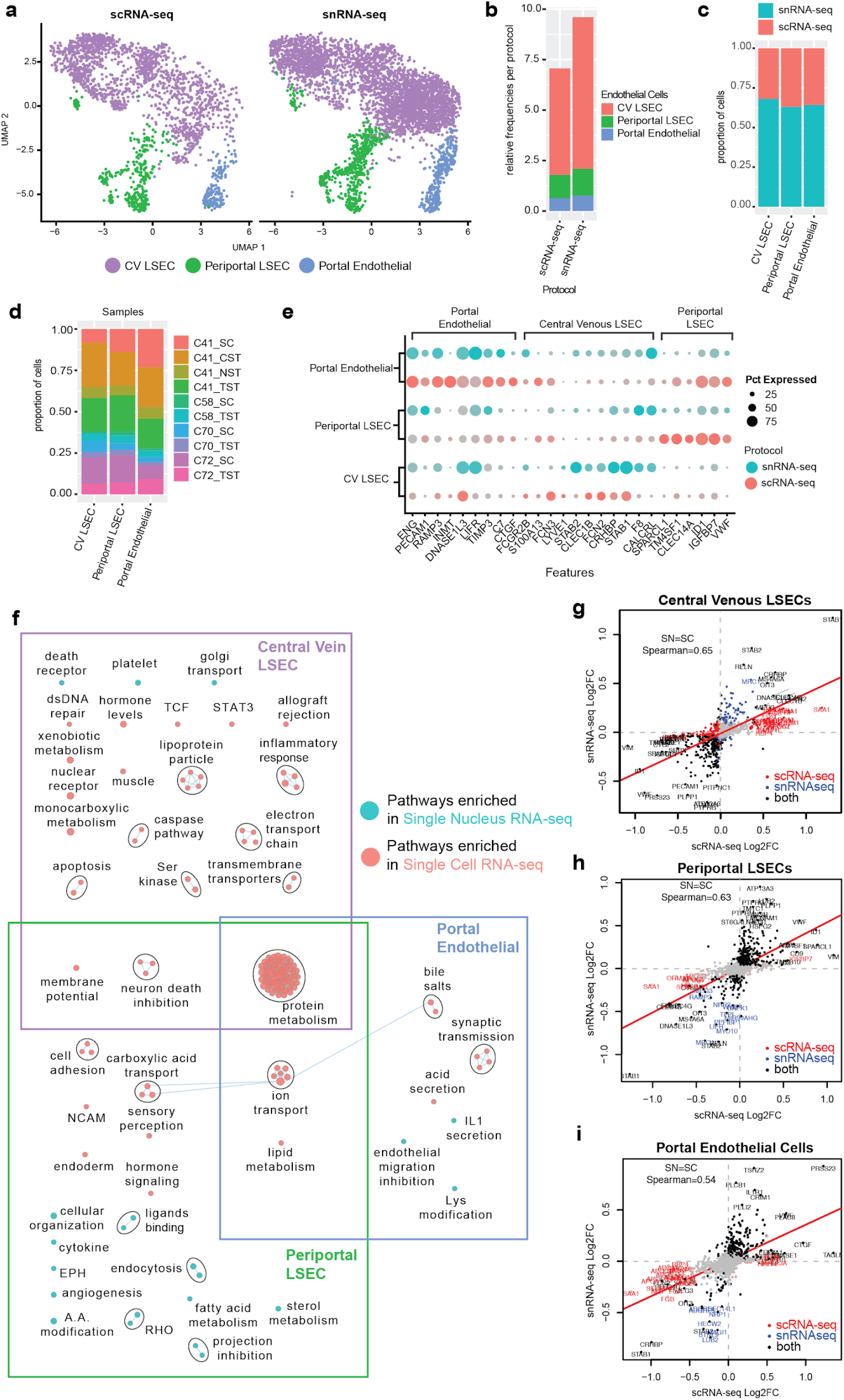
Analysis of LSECs in the combined scRNA-seq and snRNA-seq dataset. **a:** UMAP plots with the three major endothelial cell populations in the combined dataset split by protocol. **b:** Frequency of each population in either scRNA-seq or snRNA-seq datasets. Distribution of each population by protocol (**c**) and by sample (**d**) in the combined dataset. **e:** Dot-plot indicating the relative expression of known LSEC marker genes in each population by protocol. The size of the circle indicates the percentage of cells in each population expressing each gene. **f:** Pathway enrichment analysis examining which cellular pathways are better represented by snRNA-seq (cyan) and scRNA-seq (pink) in each of the LSEC subpopulations. Circles (nodes) represent pathways, sized by the number of genes included in that pathway. Related pathways, indicated by light blue lines, are grouped into a theme (black circle) and labeled. **g**,**i:** Log2FC of significant genes (q-value<0.05) within either scRNA-seq (Red) or snRNA-seq (blue) or both (black) for each cluster within the LSEC dataset. LSEC: Liver Sinusoidal Endothelial cells.

### Intrahepatic monocytes/macrophages

We previously characterized two populations of intrahepatic macrophages with immunoregulatory or inflammatory properties respectively.(2) We identified both of these populations in sc and sn RNA-seq, however we note a higher overall proportion of macrophages in snRNA-seq (7.4% of nuclei vs 4.1% of cells) and a lower relative proportion of inflammatory macrophages in snRNA-seq (46% vs 59%) (Fig. 6a-d, Supplementary Table 9). Although snRNA-seq was more efficient at capturing non-inflammatory macrophages and their associated marker genes (Fig. 6b,e), several marker genes for these populations are present in both snRNA-seq and scRNA-seq maps (*CD68, PTPRC, MARCO*). Meanwhile, the markers used to describe inflammatory macrophages (*LYZ, S100A8, S100A9*) were better represented byscRNA-seq (Fig. 6f). Immune-associated pathways : IFNg, leukocyte activation, phagocytosis, bacterial response, were more highly expressed in snRNA-seq rather than scRNA-seq (Fig. 6g), suggesting that macrophages may be dissociation-sensitive.

**Fig. 6:**
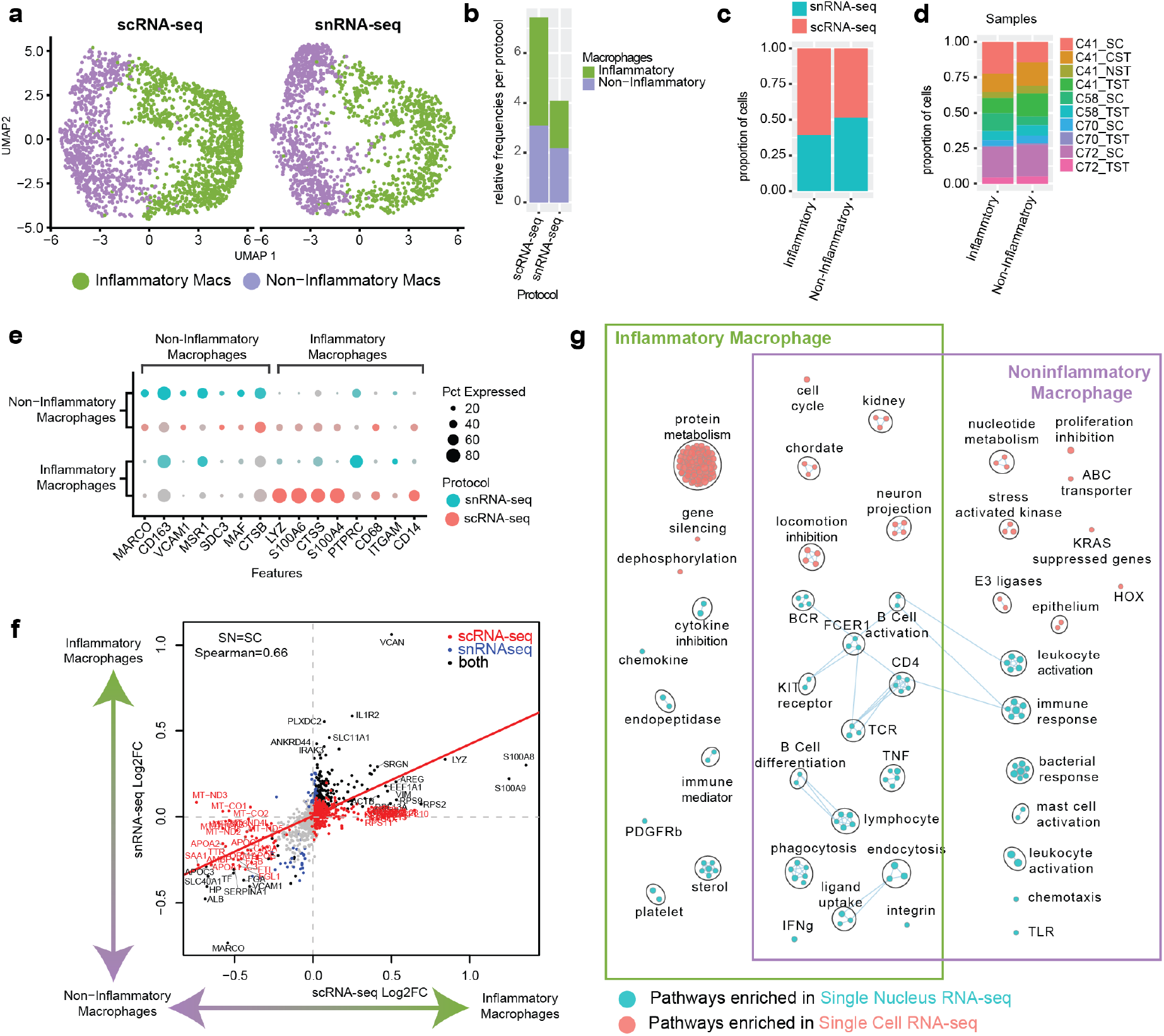
Analysis of liver resident macrophages in the combined scRNA-seq and snRNA-seq dataset. **a**: UMAP plots depicting the clustering of inflammatory and non-inflammatory macrophages in the combined dataset split by protocol. **b**: stacked bar-plot indicating the frequency of each population in either scRNA-seq or snRNA-seq datasets. Distribution of each population by protocol (**c**) and by sample (**d**) in the combined dataset. **e:** Dot-plot indicating the relative expression of known inflammatory and non-inflammatory macrophage marker genes in each cluster by protocol. The size of the circle indicates the percentage of cells in each population expressing each gene. **f:** Log2FC of significant genes (5% FDR) within either scRNA-seq (Red) or snRNA-seq (blue) or both (black) for each cluster within the macrophage populations, non-significant shown in grey. **g**: Pairwise pathway enrichment analysis comparing snRNA-seq to scRNA-seq in each macrophage subpopulation. Pathways enriched in snRNA-sq are labelled in cyan and pathways enriched in scRNA-seq are indicated in pink. Circles (nodes) represent pathways, sized by the number of genes included in that pathway. Related pathways, indicated by light blue lines, are grouped into a theme (black circle) and labeled. Macs: Macrophages.

### Intrahepatic lymphocyte populations

We previously observed that lymphocytes are well-detected after hepatic tissue disruption.(2). Major lymphocyte populations were captured by each protocol (Fig. 7a) and in each sample (Fig. 7b), lymphocytes comprise 5.2% of the scRNA-seq dataset (1524/29432) but make up only 1.8% of the snRNA-seq dataset (804/43863). All lymphocyte subpopulations were captured at higher frequency by scRNA-seq (Fig. 7c,d). B cell populations in particular make up only 0.2% of the snRNA-seq dataset but are enriched almost 2x for in scRNA-seq data (0.5%) (Supplementary Table 10). In our examination of marker genes for each cluster present across both protocols (Fig. 7e), *IL7R* and *S100A4* serve as best markers for resident memory T cells. The top marker genes for γδ T cells and NK cells have significant overlap. Unfortunately, B cell receptor and T cell receptor genes were not well-captured by snRNA-seq (Extended Data Fig. 24). As such, scRNA-seq captures transcripts that provide the resolution for differentiating distinct lymphocyte populations.

**Fig. 7:**
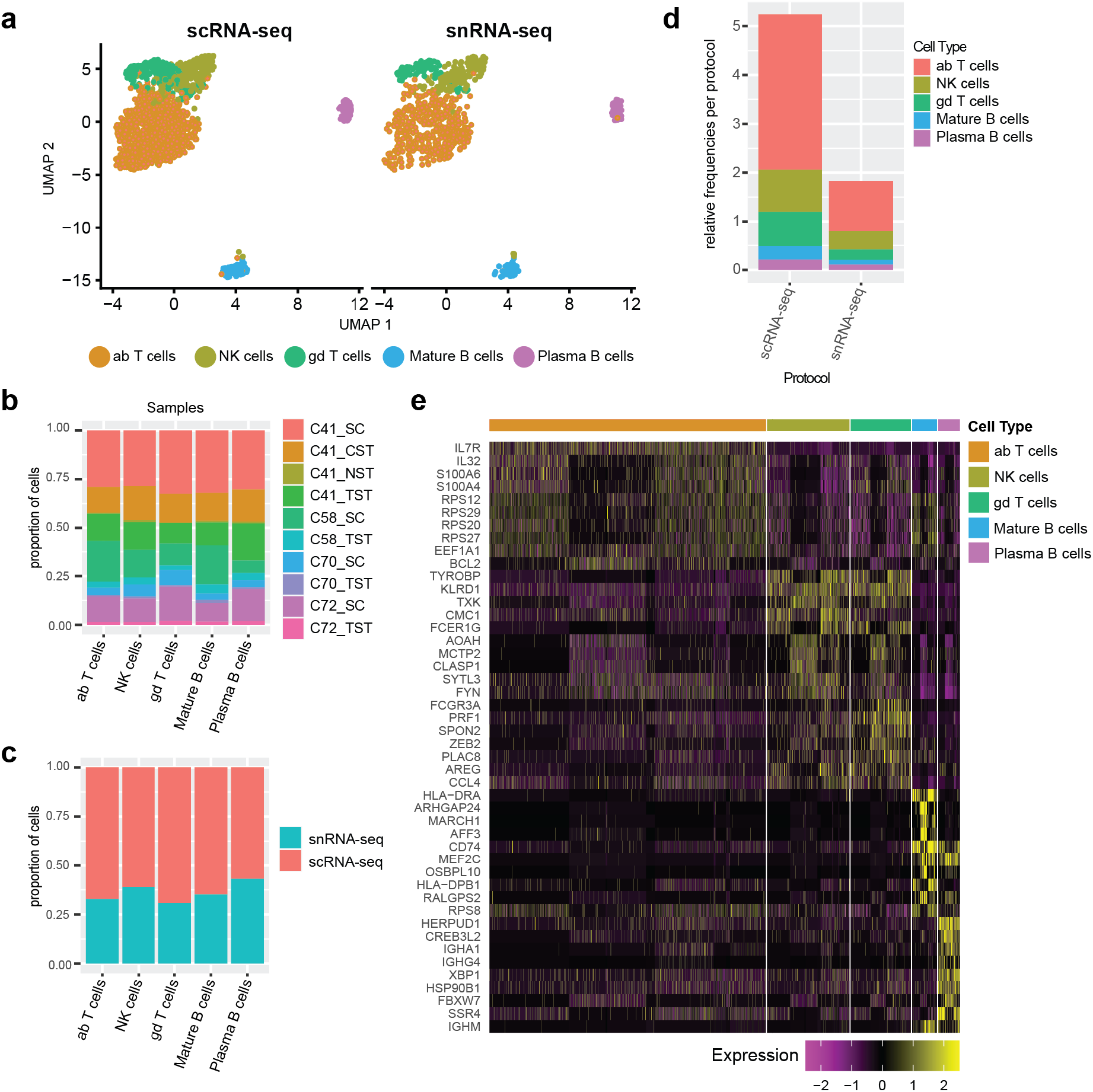
Liver-resident lymphocytes are enriched in scRNA-seq datasets. **a**: UMAP plots depicting the clustering of various lymphocyte subpopulations in the combined dataset split by protocol. **b**: Frequency of each population in either scRNA-seq or snRNA-seq datasets. Distribution of each population by protocol (**c**) and by sample (**d**) in the combined dataset. **e:** Heat-map showing the most-significantly upregulated genes per cluster. ab: alpha-beta; gd: gamma-delta; NK: natural killer.

## DISCUSSION

Comparing single-cell and single-nucleus RNA-seq protocols applied to four samples of healthy human liver, revealed several significant differences. Both techniques produced high quality data that could elucidate the major cell classifications in the human liver. However, cell-type frequencies are distorted in scRNA-seq, mainly due to the resiliency of immune cells to tissue dissociation compared to parenchymal cells. These differences impact the sensitivity of each method to delineate subtypes of the respective cell types. SnRNA-seq enables the detection of many additional cell subtypes of cholangiocytes and stellate cells which are difficult to distinguish using scRNA-seq. For example, the transcriptional profile of small duct cholangiocytes provided in this dataset may act as a reference for assessing how cholangiocytes in small-duct cholangio-pathologies like primary biliary cholangitis (PBC) might differ transcriptionally to those in the healthy liver. Furthermore, this platform could potentially allow for distinguishing differences in the cellular landscape of poorly understood autoimmune liver diseases, for example differentiating between PBC and primary sclerosing cholangitis, which is a disease mainly affecting large bile ducts.

A complete characterization of the intrahepatic immune landscape is also crucial to understanding the pathogenesis of liver disease. Despite the advantages of nuclei profiling, many important markers of immune cells were completely absent from snRNA-seq data. For instance, none of the T cell receptor or B cell receptor components were detected in our single-nucleus RNA-seq samples. Thus, we recommend that studies investigating intrahepatic immune populations use scRNA-seq.

The ability of snRNA-seq to overcome the limitations associated with tissue dissociation and to capture parenchymal cells in high resolution, opens a new avenue for a detailed examination of the interplay between parenchymal and NPCs in health and disease. Additionally, the ability to perform snRNA-seq in frozen tissues can enable the examination of biobanked samples with single-cell resolution. Taken together, we have shown that single-cell and single-nucleus RNA-seq generate high-quality data from normal liver samples, with single nucleus allowing for better examination of parenchymal cells, including stellate cells and cholangiocytes. This combined dataset enables a complex examination of parenchymal cells complexity and provides a foundation for single cell liver disease studies.

## EXPERIMENTAL PROCEDURES

### Preparation of fresh tissue homogenates and nuclei from snap-frozen human liver tissue

Human liver tissue from the caudate lobe was obtained from neurologically-deceased donor liver acceptable for liver transplantation. Samples were collected with institutional ethics approval from the University Health Network (REB# 14-7425-AE). A 3mm-cubed fragment of tissue was preserved for snRNA-seq by snap freezing in liquid nitrogen. Single cell suspensions of fresh human liver were generated as previously described (2) two step collagenase perfusion protocol [https://doi.org/10.17504/protocols.io.m9sc96e]. Single nucleus extraction was performed as previously described (9). The full description of processing is found in the extended methods.

### 10x sample processing, cDNA library preparation, sequencing and data processing

Samples were prepared as outlined by the 10x Genomics Single Cell 3′ v2 and 3’ v3 Reagent Kit user guides and as described previously (2). Single cell data was processed using 10x Cell Ranger software version 3.01, mapping reads to the GRCh38 human genome. Single nucleus data was processed using Cell Ranger version 3, and reads were mapped to a modified transcriptome based on GRCh38 which included intronic regions to ensure quantification of reads derived from immature, unspliced mRNA present in the nucleus. The full description of 10x sample and data processing is found in the extended methods.

### Data integration and clustering

The data was integrated using default parameters of Harmony (10), then clustered using Seurat’s SNN-Louvain clustering algorithm (30). The data was clustered using 30 sets of parameters, and the most consistent clusterings were identified using apcluster (31) on the cluster-cluster distance matrix calculated using the Variation of Information criterion (32). The full description of data integration and clustering can be found in the extended methods.

### Gene-type and gene length biases between scRNA-seq and snRNA-seq data

3,804 housekeeping genes were obtained from the literature (11). Long non-coding RNAs and protein coding gene lists were obtained from Ensembl. Nuclear-encoded mitochondrial proteins were obtained from MitoCarta3.0 (33). Ribosomal genes were obtained from the Ribosomal Protein Gene database (34). Transcript GC content, miRNA binding sites, transcript length, UTR lengths and intron length were obtained from Ensembl Biomart. Log Fold changes of mean expression across all cell-types in scRNA-seq and snRNA-seq were calculated across all samples.

### Pathway enrichment, correlation and trajectory inference analysis

Pathway enrichment analysis was performed as previously described (2) with the addition of a dissociation signature to the pathway gene set database (16). Slingshot (v1.8.0) was employed to infer the pseudotime based on the Harmony embedding matrix of cells. Lineages were calculated using the Slingshot UMAP embedding protocol (18). Diffusion maps (35) (destiny, v3.1.1) were computed with both the raw counts matrix and the PCA loadings. Spearman’s rank correlation coefficient was calculated on each pair of outputs of these analyses and plotted using corrplot (v0.84). The full description of this analysis is found in the extended methods.

### Validation of zonated gene signatures using spatial transcriptomics

Healthy human liver tissue was embedded in OCT, frozen and cryosectioned with 16um thickness at −10C (cryostar NX70 HOMP). Sections were placed on a chilled Visium Tissue Optimization Slide (10x Genomics) and processed following the Visium Spatial Gene Expression User Guide. Tissue was permeabilized for 12 minutes, based on an initial optimizations trial and libraries were prepared according to the Visium Spatial Gene Expression User Guide. Samples were sequenced on a NovaSeq 6000.

### Visium spatial transcriptomics

The Visium spatial transcriptomic data was sequenced to a depth of 167,400,637 reads, a saturation of 77%. These reads were mapped to the GRCh38 human genome and expression was quantified with the spaceranger-1.1.0. Further processing and visualization was performed with Seurat (version 3.2.1). The full description of Visium data processing is found in the extended methods.

### Validation of zonated protein expression *via* the Human Protein Atlas

Immunostaining images were obtained from the Human Protein Atlas (https://www.proteinatlas.org) (19). Lobule annotation was confirmed by a liver pathologist (C. Thoeni).

## Abbreviations

scRNA: seq-single cell RNA sequencing
snRNA: seq-single nucleus RNA sequencing
GSEA: gene set enrichment analysis
LSECs: liver sinusoidal endothelial cells
HSC: Hepatic Stellate cells

## Data availability

Raw data, processed data, code and all cluster-specific DE genes will be made publicly available through respective databases upon acceptance.

## Acknowledgements

This project has been made possible in part by grant number CZF2019-002429 from the Chan Zuckerberg Initiative DAF, an advised fund of Silicon Valley Community Foundation. This research was supported in part by the University of Toronto’s Medicine by Design initiative, which receives funding from the Canada First Research Excellence Fund (CFREF) to SAM, GDB and IDM; by the NRNB (U.S. National Institutes of Health, grant P41 GM103504) to GDB and by the Toronto General and Western Hospital Foundation. JA has received graduate fellowships from NSERC (CGS-M) and an Ontario Graduate Scholarship. CTP has received postdoctoral funds from the Canadian Network on Hepatitis C (CanHepC) and PSC Partners Canada. CanHepC is funded by a joint initiative of the Canadian Institutes of Health Research (CIHR) (NHC-142832) and the Public Health Agency of Canada (PHAC).

