## Supplementary Material for "Single Cell, Single Nucleus and Spatial RNA Sequencing of the Human Liver Identifies Hepatic Stellate Cell and Cholangiocyte Heterogeneity"

**List of Extended Data Figures** **Page**

Extended Data Fig. 1: Background contamination of hepatocytes in other cell types (estimates per cell). 3

Extended Data Fig. 2: Uncorrected and globally scaled data: Cells cluster by technology rather than cell type. 4

Extended Data Fig. 3: Log fold change in UMI expression as detected in snRNA-seq relative to single cell RNA-seq broken down by genomic characteristics. 5

Extended Data Fig. 4: Log fold change in UMI expression as detected in single nucleus RNA-seq relative to single cell RNA-seq is related to gene length. 7

Extended Data Fig. 5: Cell-types are distributed across samples. 8

Extended Data Fig. 6: Cells and Nuclei subsets comprising each core cell type that were selected for further examination. 9

Extended Data Fig. 7: Validation of human hepatocyte cluster annotation with bulk RNA-seq. 10

Extended Data Fig. 8: Spatial distribution of Landmark pericentral and periportal hepatocyte markers across the healthy human liver using spatial transcriptomics (10x genomics Visium platform). 11

Extended Data Fig. 9: Validation of human central venous hepatocyte cluster by Immunostaining. 13

Extended Data Fig. 10: Validation of human periportal hepatocyte cluster by Immunostaining. 14

Extended Data Fig. 11: Spatial distribution of hepatocyte cluster associated gene signature and the top differentially expressed genes from interzonal-like clusters across the healthy human liver using spatial transcriptomics (10x genomic Visium platform). 15

Extended Data Fig. 12: Validation of human interzonal hepatocyte cluster by Immunostaining. 16

Extended Data Fig. 13: The expression of common cholangiocyte, stem cell and hepatocyte markers in the cholangiocyte sub-clusters. 17

Extended Data Fig. 14: Bile duct enrichment of cholangiocyte and BCL2+ cholangiocyte specific marker proteins via Immunohistochemistry 19
Extended Data Fig. 15: Breakdown of pathways enriched in scRNA-seq vs snRNA-seq in the cholangiocyte sub-populations. 20
Extended Data Fig. 16: Differences in upregulated genes between scRNA-seq and snRNA-seq for cholangiocyte subpopulations. 22
Extended Data Fig. 17: Pathway enrichment analysis examining active cellular pathways in the cholangiocyte-associated subpopulations. 23
Extended Data Fig. 18: Progenitor-associated markers Immunostaining from the Human Protein Atlas. 24
Extended Data Fig. 19: Spatial distribution of cholangiocyte-associated subpopulations across the healthy human liver using spatial transcriptomics (10x genomic Visium platform). 25
Extended Data Fig. 20: Cholangiocyte trajectory inference. 26
Extended Data Fig. 21: Hepatic stellate cell markers and trajectory inference. 27
Extended Data Fig. 22: Hepatic stellate cell genes and gene signatures in the healthy human liver using spatial transcriptomics (10x genomic Visium platform). 29
Extended Data Fig. 23: Verification of macrophage clusters in the healthy human liver using spatial transcriptomics (10x genomic Visium platform). 30
Extended Data Fig. 24: Expression of TCR and BCR components in scRNA-seq and snRNA-seq. 31

**List of Supplementary Tables** **Page**

Supplementary Table 1: Core cell annotation markers 32
Supplementary Table 2: RNA composition of scRNA and snRNA-seq - mean±SD 32
Supplementary Table 3: Median UMI counts and number of detected genes per cell / nucleus across all samples within each sample. 32
Supplementary Table 4: Cell Type Compositions for scRNA-seq and snRNA-seq. 33
Supplementary Table 5: Frequency of Hepatocyte clusters in scRNA-seq and snRNA-seq 33
Supplementary Table 6: Representation of cholangiocyte clusters in scRNA/ snRNA-seq 34
Supplementary Table 7: Hepatic Stellate cell distribution 35
Supplementary Table 8: Further Description of LSEC Populations. 35
Supplementary Table 9: Further Description of Macrophages 35
Supplementary Table 10: Further Description of Lymphocyte populations. 36
Supplementary Table 11: cellranger summary 36
Supplementary Table 12: SOUPX genes: 36
Extended Experimental Procedures 36

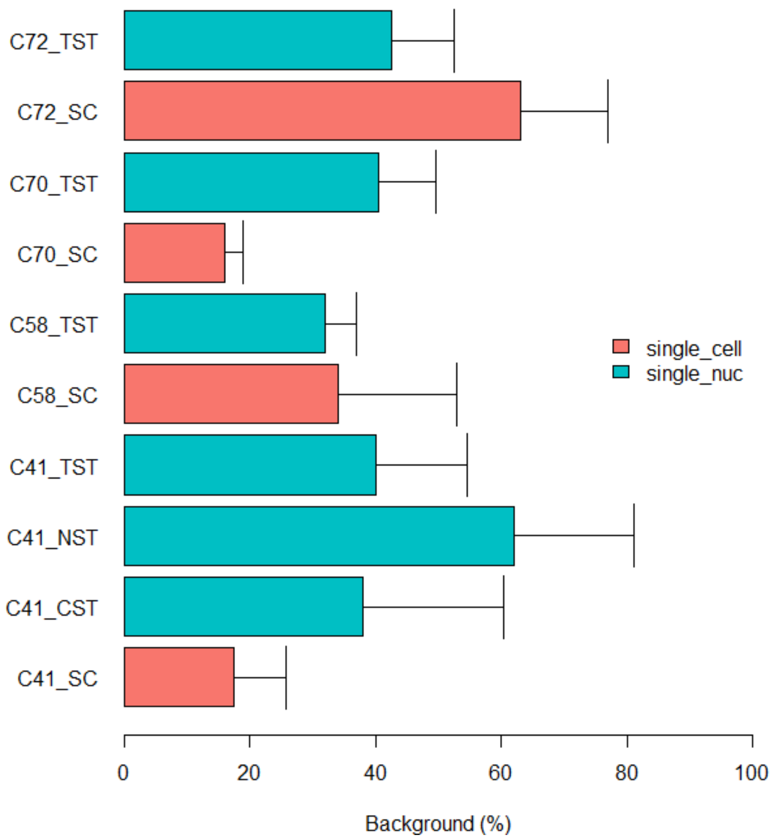


**Extended Data Fig. 1: Background contamination of hepatocytes in other cell types (estimates per cell).** This estimates how much of the total library comes from ambient RNA. This percentage is calculated by looking for lists of genes that are not expected to be expressed in each cluster using SoupX (1). Error bars indicate standard deviations of the mean since the large number of cells reduces standard errors to be indistinguishable from the bars.

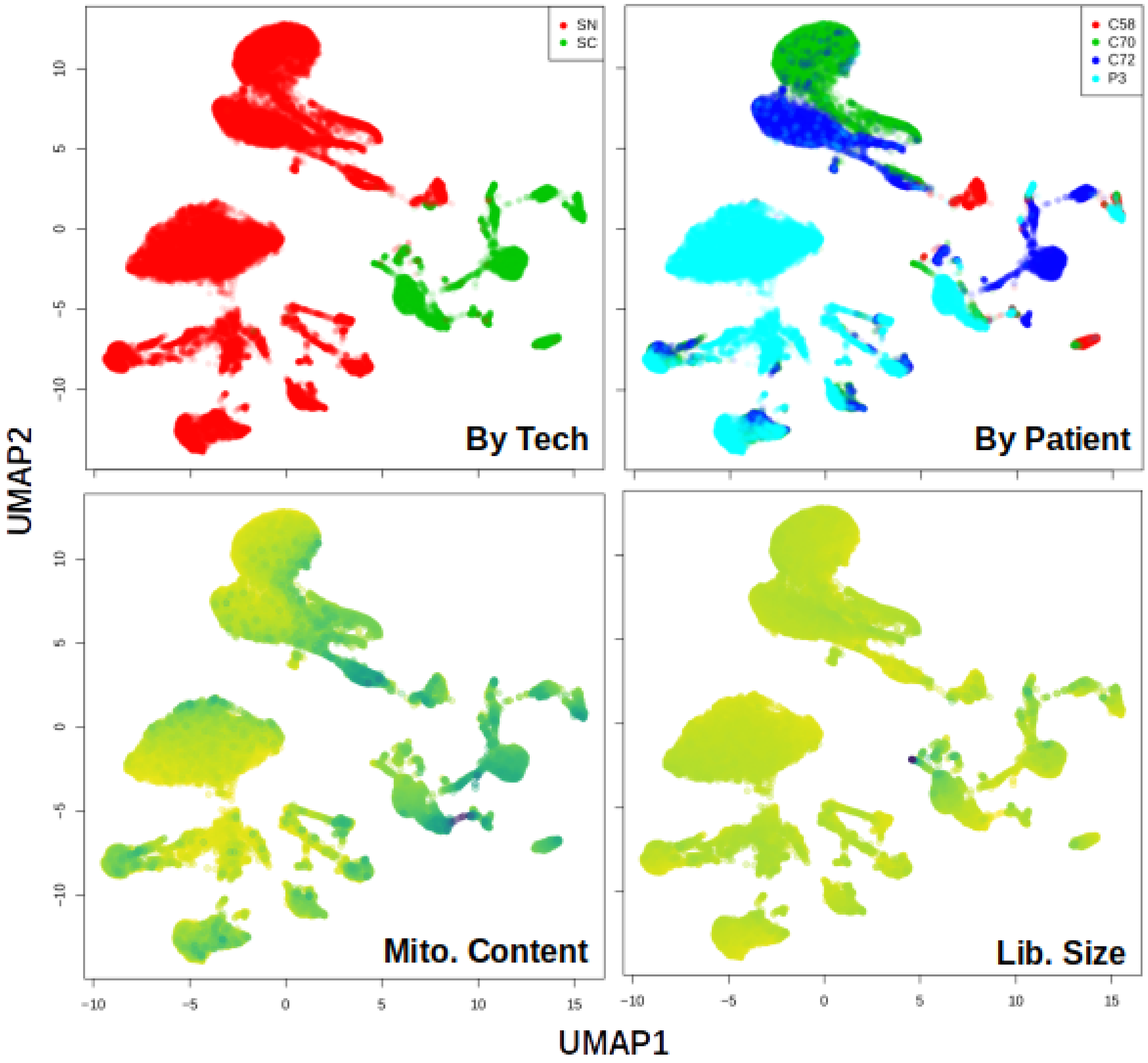


**Extended Data Fig. 2: Uncorrected and globally scaled data: Cells cluster by technology and patients rather than cell type.** UMAP plots where cells are coloured by technology (top-left), by patient (top -right), average mitochondrial gene proportion (bottom-left) and by library size (bottom right).

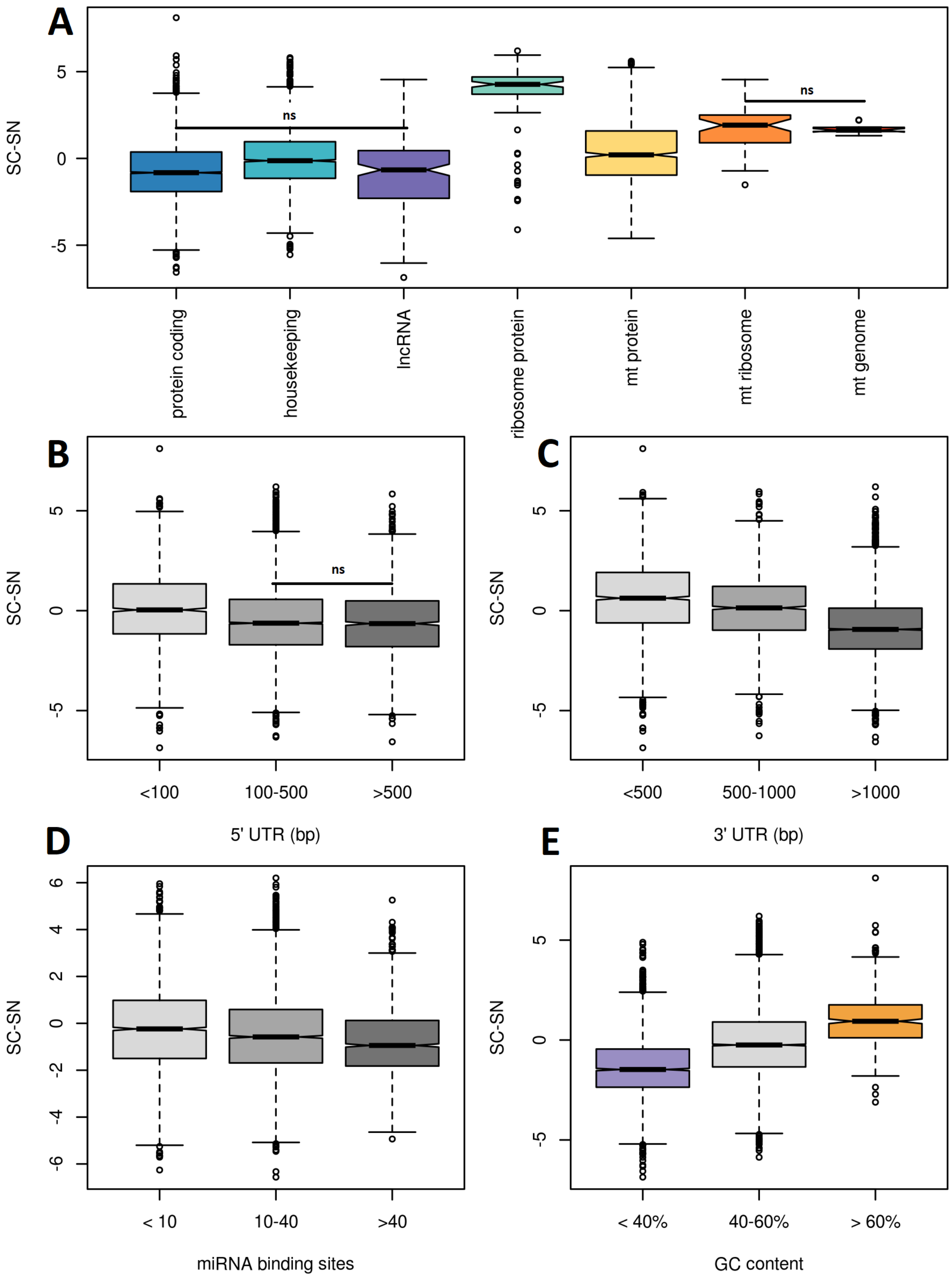

A
SC-SN
5
0
-5
ns
ns
protein coding
housekeeping
lncRNA
ribosome protein
mt protein
mt ribosome
mt genome
B
SC-SN
ns
<100
100-500
>500
5' UTR (bp)
C
SC-SN
<500
500-1000
>1000
3' UTR (bp)
D
SC-SN
6
4
2
0
-2
-4
-6
< 10
10-40
>40
miRNA binding sites
E
SC-SN
< 40%
40-60%
> 60%
GC content

**Extended Data Fig. 3: Log fold change in UMI expression as detected in snRNA-seq relative to single cell RNA-seq broken down by genomic characteristics.** a, By gene biotype whether genes are housekeeping genes or not, ribosomal and mitochondrial proteins are higher in single cell RNA-seq while non-housekeeping protein coding genes and lncRNAs are higher in single nucleus RNA-seq. b-c, Genes with longer UTRs are higher in single nucleus RNA-seq. d, genes with more microRNA binding sites are higher in single nucleus. e, genes with lower GC are higher in single nucleus. Notches indicate 95% confidence intervals of the median, all differences except those indicated were significant at $p < 10^{-5}$ using a Wilcoxon-rank-sum test. In contrast to previous reports (2,3) long noncoding RNAs were not over-represented in snRNA-seq compared to other tissue or cell-type specific protein-coding RNAs.

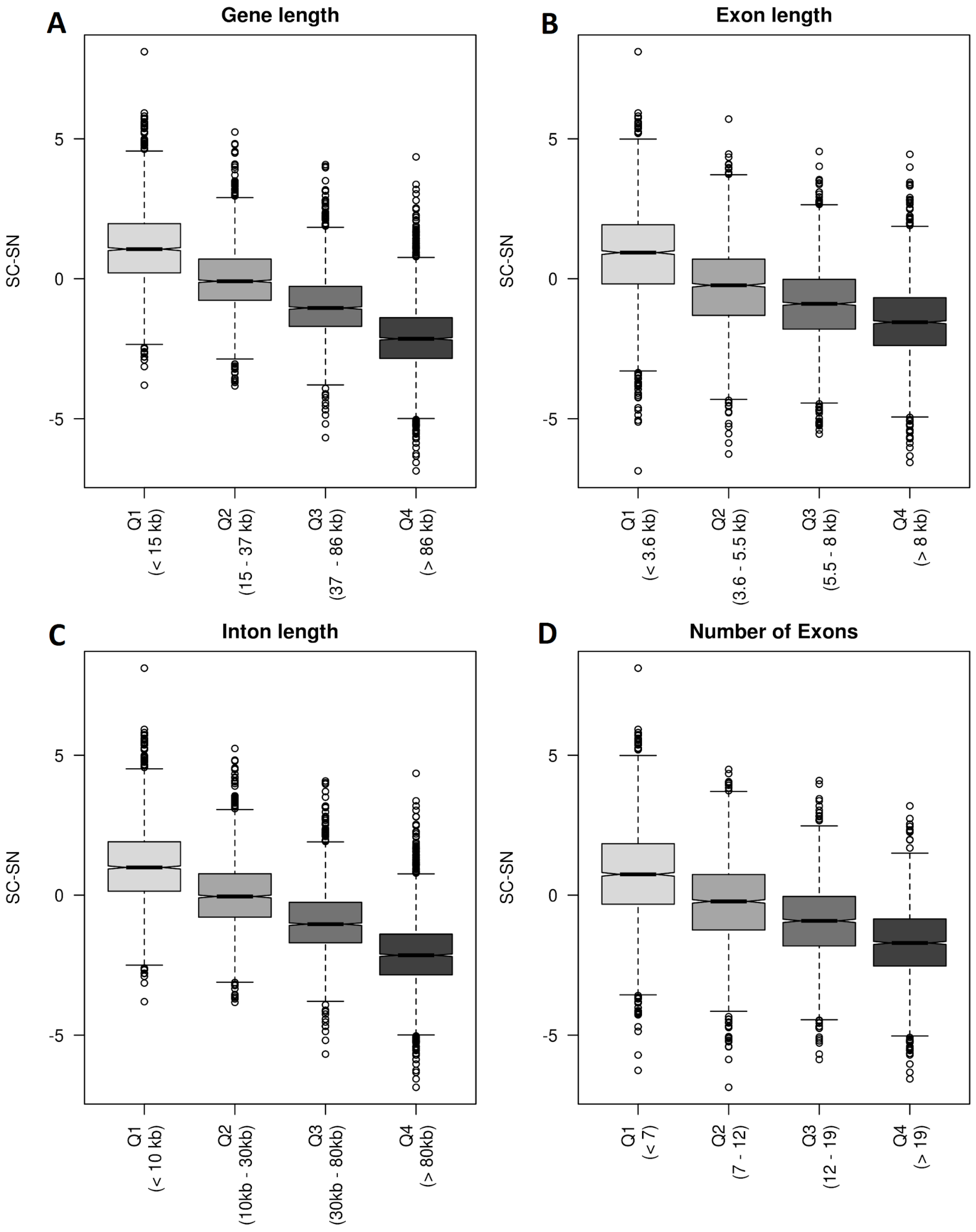


**Extended Data Fig. 4: Log fold change in UMI expression as detected in single nucleus RNA-seq relative to single cell RNA-seq is related to gene length.**

All transcripts were aggregated to calculate total lengths. (a) Total gene length including UTRs, introns, and exons. (b) Total exon length. (c) Total intron length (d) number of exons. Notches indicate 95% confidence intervals of the median, all differences were significant at $p < 10^{-10}$ using a Wilcoxon-rank-sum test. In agreement with prior data, (4) we note biased expression of long genes in snRNA-seq, particularly genes with large introns.

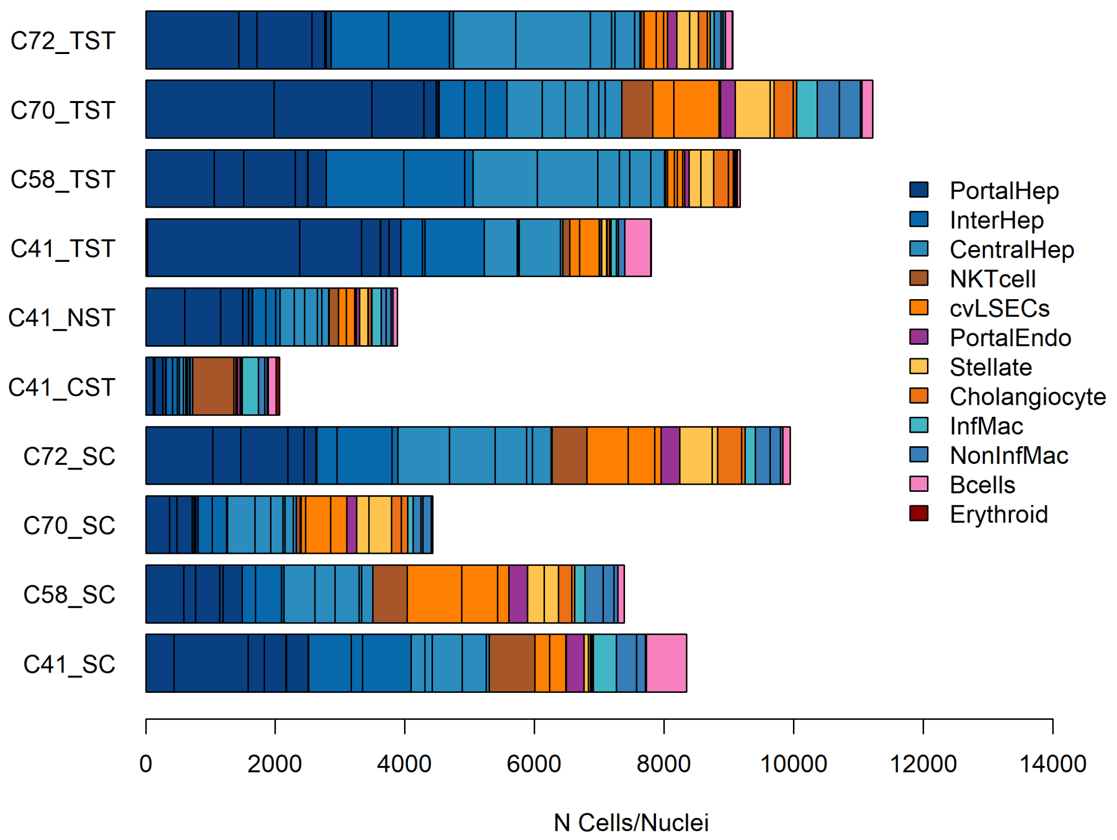


**Extended Data Fig. 5: Cell-types are distributed across samples.** The number of cells or nuclei identified in each of the 34 clusters is shown, dark lines separate the counts for each cluster and clusters are coloured based on the manually annotated cell-types. C41-C72 identifies the individual the sample originated from while CST, NST, TST are three different single nuclei extraction detergents, and SC is single-cell RNA-seq.

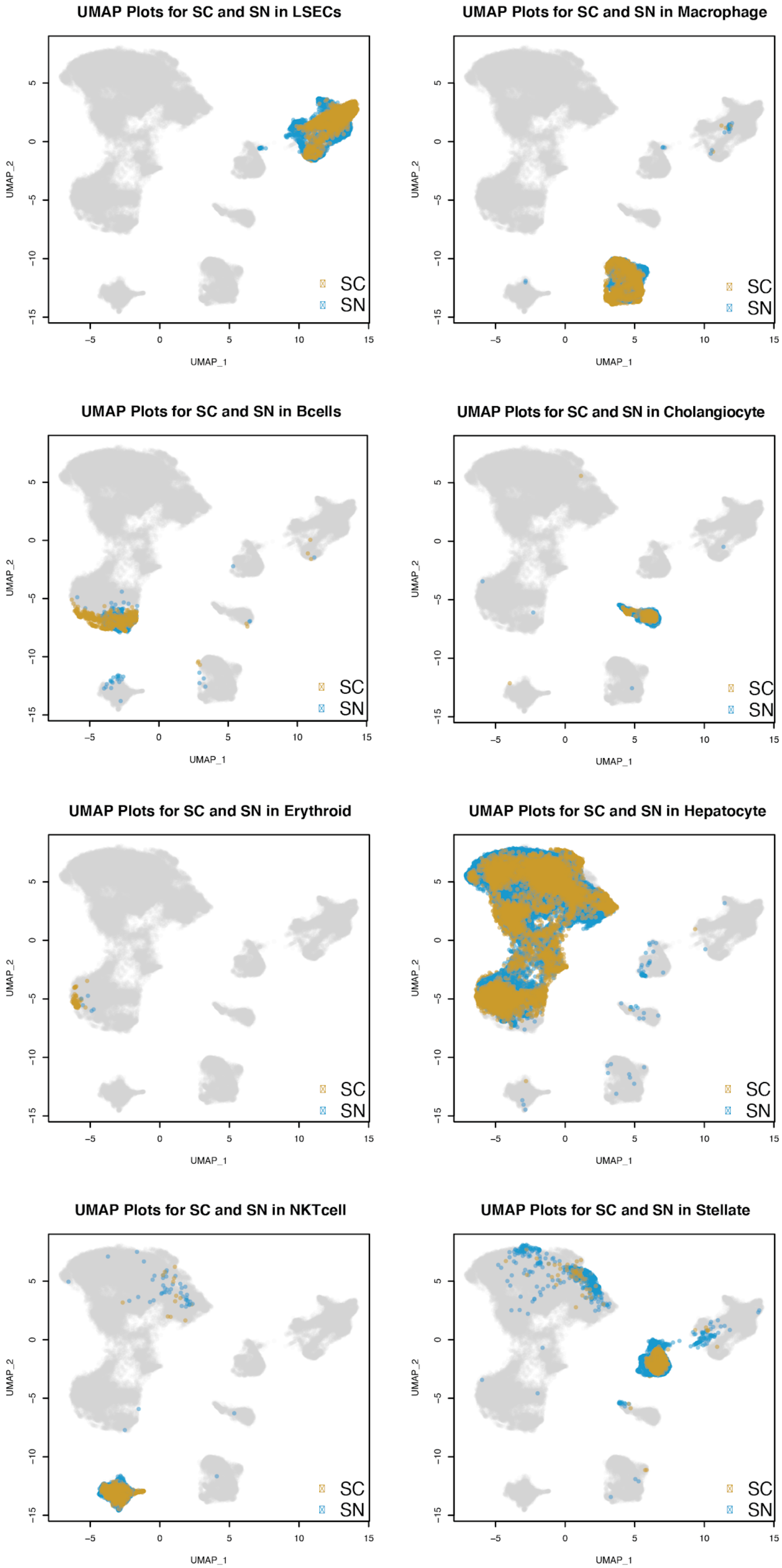
UMAP Plots for SC and SN in LSECs
UMAP Plots for SC and SN in Macrophage
UMAP Plots for SC and SN in Bcells
UMAP Plots for SC and SN in Cholangiocyte
UMAP Plots for SC and SN in Erythroid
UMAP Plots for SC and SN in Hepatocyte
UMAP Plots for SC and SN in NKTcell
UMAP Plots for SC and SN in Stellate
SC
SN
UMAP_1
UMAP_2

**Extended Data Fig. 6: Cells and nuclei subsets comprising each core cell type that were selected for further examination.**

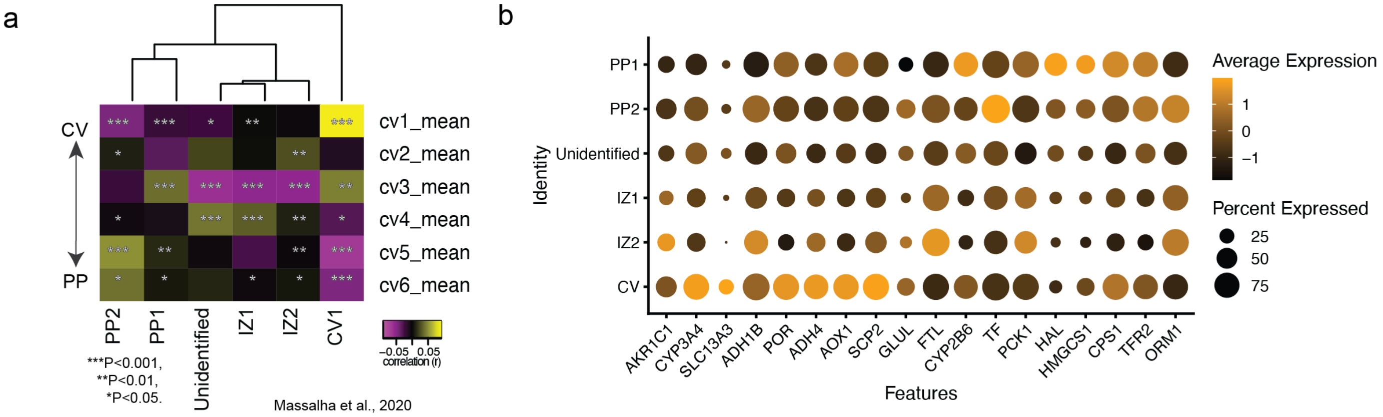


**Extended Data Fig. 7: Validation of human hepatocyte cluster annotation with bulk RNA-seq.** a, Correlation of human hepatocyte clusters to laser capture microscopy linked bulk RNA-seq of six human sinusoid layers calculated using Spearman correlation (5). ****P*<0.001, ***P*< 0.01, **P*<0.05. b, Dot-plot of relative expression levels of differentially upregulated genes in central venous and periportal hepatocyte associated genes in the combined scRNA-seq and snRNA-seq dataset.

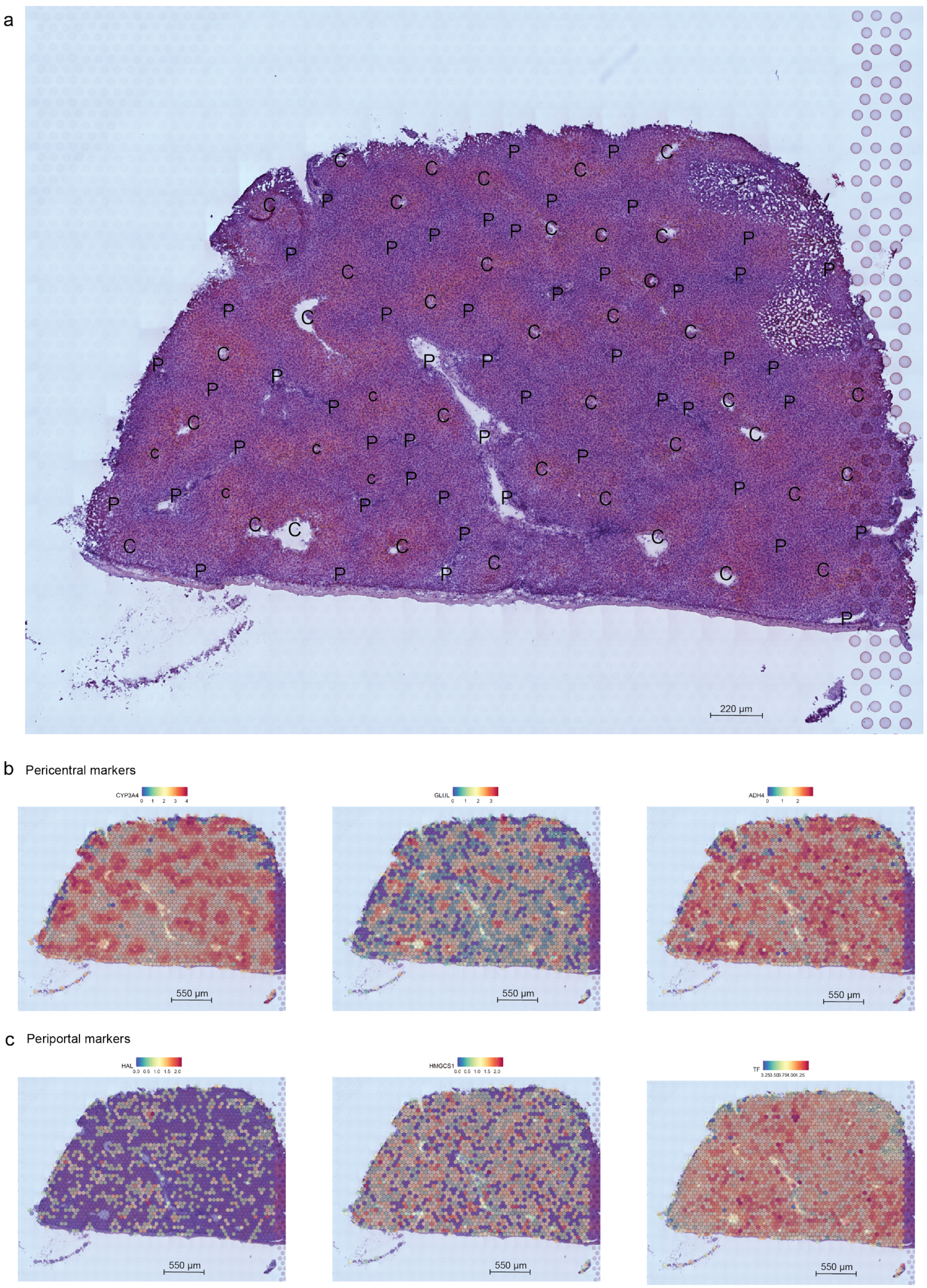
a
220 µm
b
Pericentral markers
CYP3A4
GLUL
ADH4
550 µm
c
Periportal markers
HAL
HMGCS1
TF
550 µm

**Extended Data Fig. 8: Spatial distribution of landmark pericentral and periportal hepatocyte markers across the healthy human liver using spatial transcriptomics (10x genomics Visium platform).** a, Hematoxylin and Eosin staining of healthy human liver cryosection (C73 caudate lobe) on the spatial transcriptomics slide. Pericentral (P) and Periportal regions are indicated. Spatial distribution of common pericentral (b, *CYP3A4, GLUL, ADH4*) and periportal markers (c, *HAL, HMGCS1, TF*).

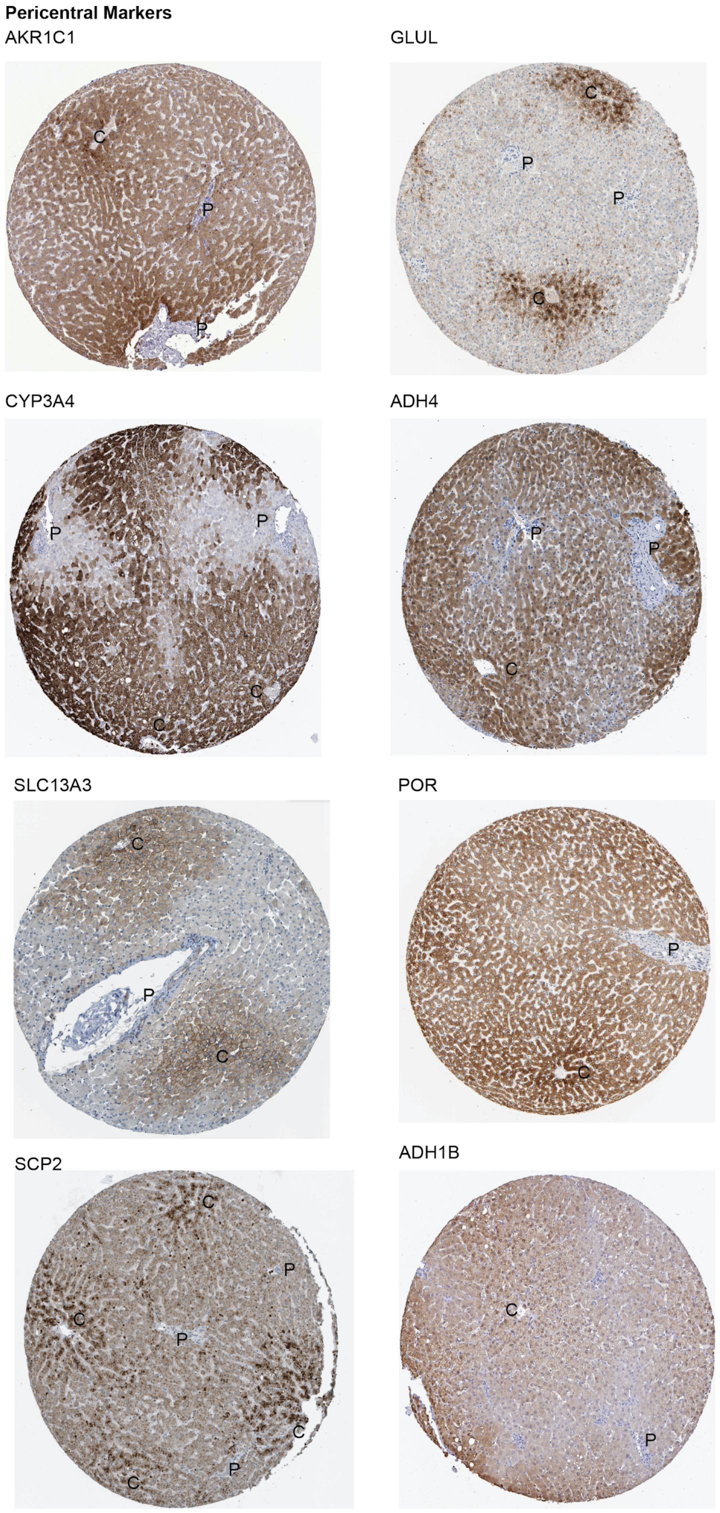


**Extended Data Fig. 9: Validation of human central venous hepatocyte cluster by Immunostaining.** Immunohistochemistry of genes differentially upregulated in central venous hepatocyte populations (CV) in the combined dataset from the Human Protein Atlas (6). Periportal (P) and Central venous (C ) regions are labeled.

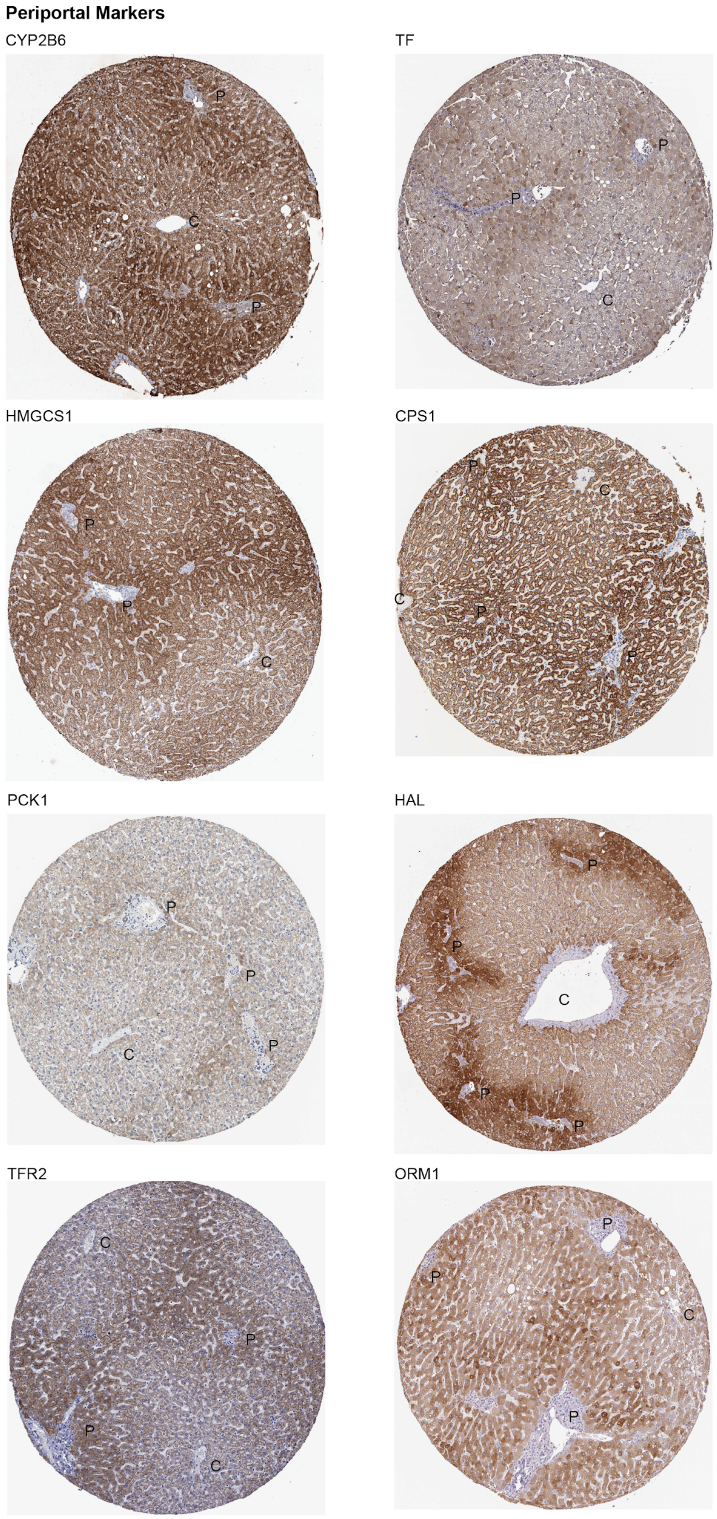


**Extended Data Fig. 10: Validation of human periportal hepatocyte cluster by Immunostaining.** Immunohistochemistry of genes differentially expressed in periportal Hepatocyte populations (P1 and P2) in the combined dataset from the Human Protein Atlas (6). Periportal (P) and Central venous (C ) regions are labeled.

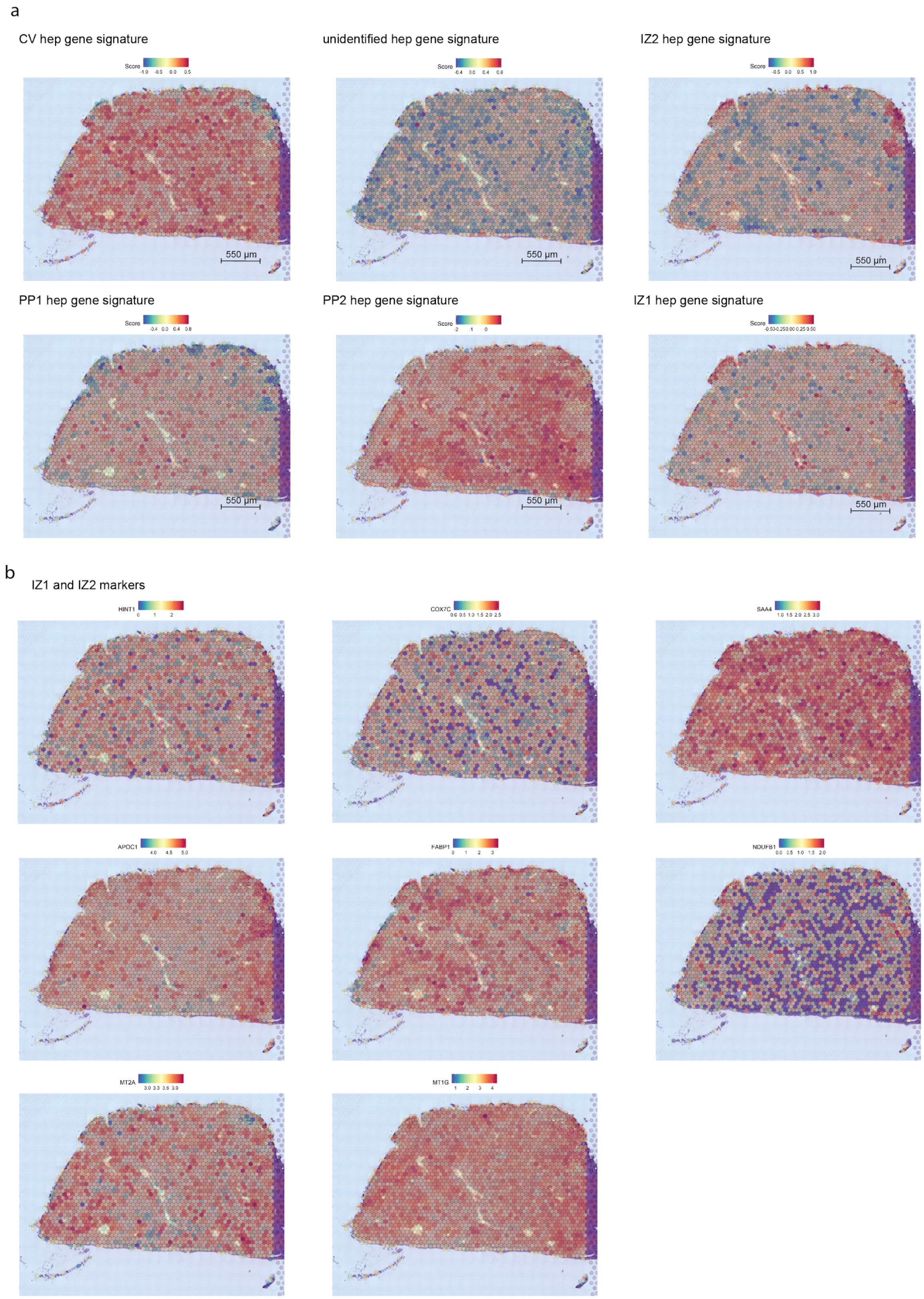
a
CV hep gene signature
unidentified hep gene signature
IZ2 hep gene signature
PP1 hep gene signature
PP2 hep gene signature
IZ1 hep gene signature
Score
550 µm
b
IZ1 and IZ2 markers
HINT1
COX7C
SAA4
APOC1
FABP1
NDUFB1
MT2A
MT1G

**Extended Data Fig. 11: Spatial distribution of hepatocyte cluster associated gene signature and the top differentially expressed genes from interzonal-like clusters across the healthy human liver using spatial transcriptomics (10x genomic Visium platform).** a, Distribution of the gene signature for each hepatocyte sub-population (Avg expression of the top 30 genes for each cluster). b, Expression of genes that are differentially expressed in the interzonal clusters in the combined scRNA-seq and snRNA-seq dataset.

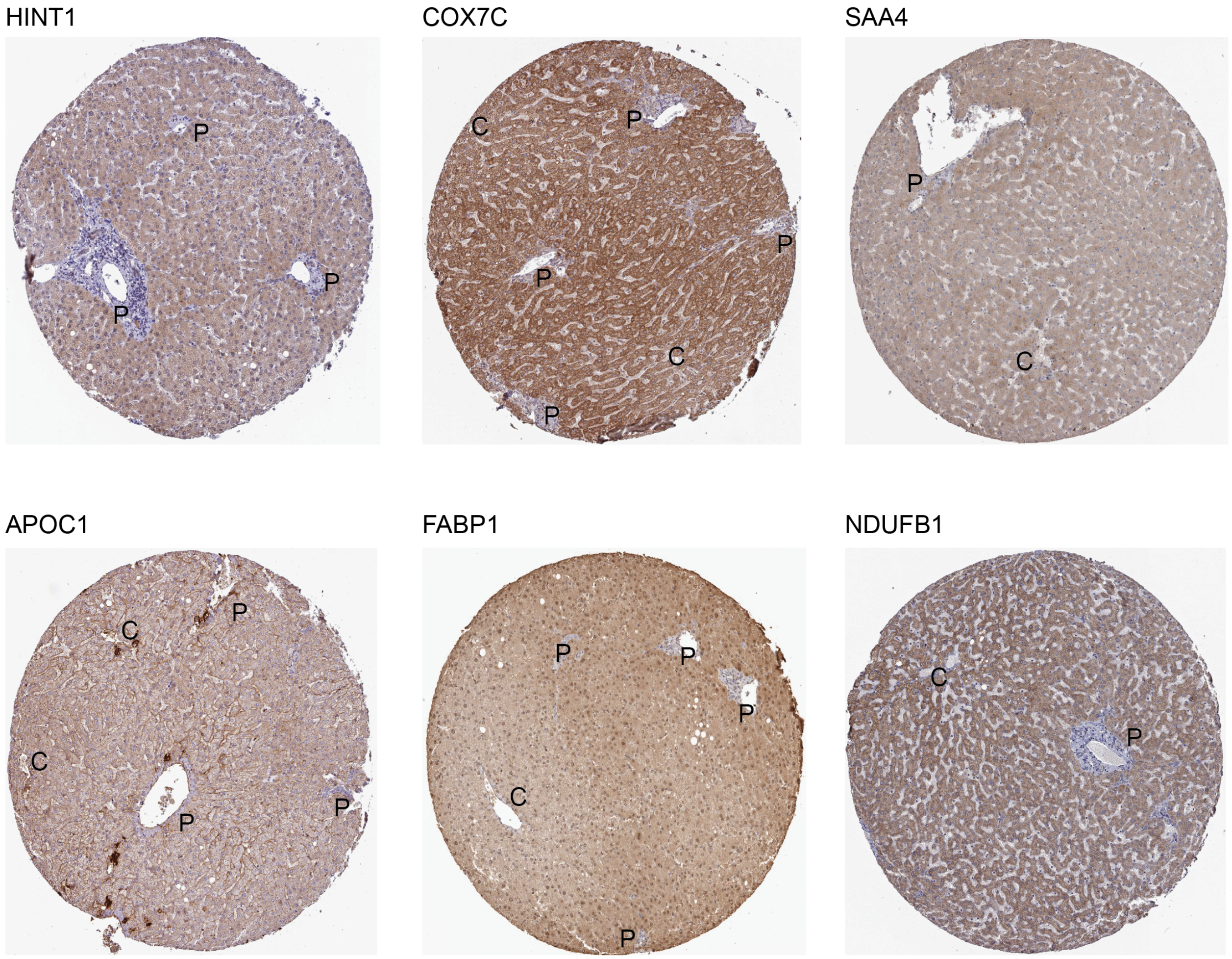


**Extended Data Fig. 12: Validation of human interzonal hepatocyte cluster by Immunostaining.** Immunohistochemistry of genes differentially upregulated in the interzonal hepatocyte population (IZ2) in the combined dataset from the Human Protein Atlas (6). C and D Periportal (P) and Central venous (C ) regions are labeled.

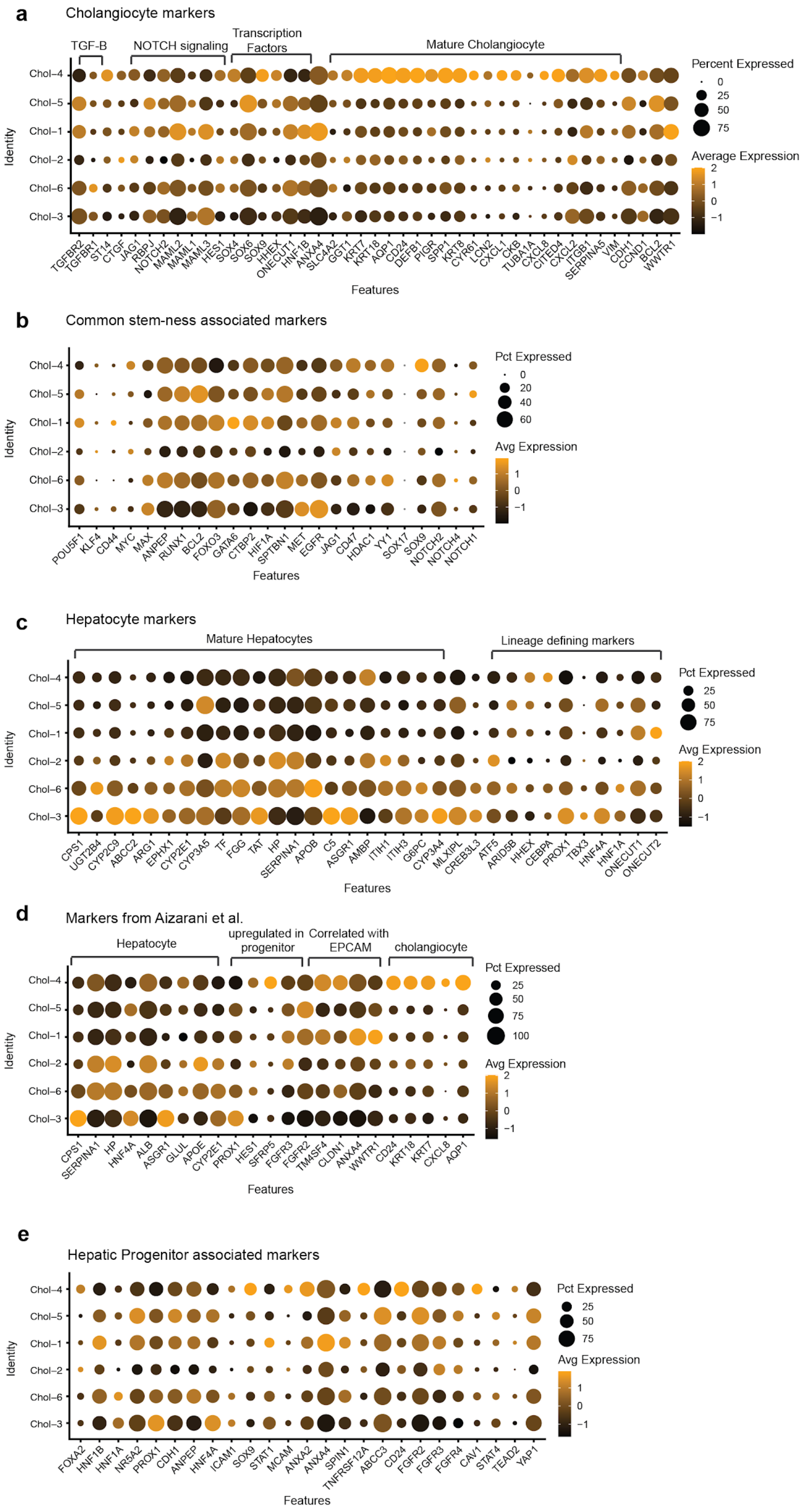
a
Cholangiocyte markers
TGF-B
NOTCH signaling
Transcription Factors
Mature Cholangiocyte
Chol–4
Chol–5
Chol–1
Chol–2
Chol–6
Chol–3
Identity
TGFBR2 TGFBR1 ST14 CTGF JAG1 RBPJ NOTCH2 MAML2 MAML1 MAML3 HES1 SOX4 SOX6 SOX9 HHEX ONECUT1 HNF1B ANXA4 SLC4A2 GGT1 KRT7 KRT18 AQP1 CD24 DEFB1 PIGR SPP1 KRT8 CYR61 LCN2 CXCL1 CKB TUBA1A CXCL8 CITED4 CXCL2 ITGB1 SERPINA5 VIM CDH1 CCND1 BCL2 WWTR1
Features
Percent Expressed
0
25
50
75
Average Expression
2
1
0
−1
b
Common stem-ness associated markers
Chol–4
Chol–5
Chol–1
Chol–2
Chol–6
Chol–3
Identity
POU5F1 KLF4 CD44 MYC MAX ANPEP RUNX1 BCL2 FOXO3 GATA6 CTBP2 HIF1A SPTBN1 MET EGFR JAG1 CD47 HDAC1 YY1 SOX17 SOX9 NOTCH2 NOTCH4 NOTCH1
Features
Pct Expressed
0
20
40
60
Avg Expression
1
0
−1
c
Hepatocyte markers
Mature Hepatocytes
Lineage defining markers
Chol–4
Chol–5
Chol–1
Chol–2
Chol–6
Chol–3
Identity
CPS1 UGT2B4 CYP2C9 ABCC2 ARG1 EPHX1 CYP2E1 CYP3A5 TF FGG TAT HP SERPINA1 APOB C5 ASGR1 AMBP ITIH1 ITIH3 G6PC CYP3A4 MLXIPL CREB3L3 ATF5 ARID5B HHEX CEBPA PROX1 TBX3 HNF4A HNF1A ONECUT1 ONECUT2
Features
Pct Expressed
25
50
75
Avg Expression
2
1
0
−1
d
Markers from Aizarani et al.
Hepatocyte
upregulated in progenitor
Correlated with EPCAM
cholangiocyte
Chol–4
Chol–5
Chol–1
Chol–2
Chol–6
Chol–3
Identity
CPS1 SERPINA1 HP HNF4A ALB ASGR1 GLUL APOE CYP2E1 PROX1 HES1 SFRP5 FGFR3 FGFR2 TM4SF4 CLDN1 ANXA4 WWTR1 CD24 KRT18 KRT7 CXCL8 AQP1
Features
Pct Expressed
25
50
75
100
Avg Expression
2
1
0
−1
e
Hepatic Progenitor associated markers
Chol–4
Chol–5
Chol–1
Chol–2
Chol–6
Chol–3
Identity
FOXA2 HNF1B HNF1A NR5A2 PROX1 CDH1 ANPEP HNF4A ICAM1 SOX9 STAT1 MCAM ANXA2 ANXA4 SPIN1 TNFRSF12A ABCC3 CD24 FGFR2 FGFR3 FGFR4 CAV1 STAT4 TEAD2 YAP1
Features
Pct Expressed
25
50
75
Avg Expression
1
0
−1

**Extended Data Fig. 13: The expression of common cholangiocyte, stem cell and hepatocyte markers in the cholangiocyte sub-clusters.** Dot-plots indicate the relative average expression of reported cholangiocyte (a), stem-cell (b), and hepatocyte (c) markers from the literature in the cholangiocyte sub-clusters in the combined scRNA-seq and snRNA-seq dataset. d, The relative average expression of cholangiocyte, hepatocyte and bipotent-progenitor marker genes from Aizarani et al. across the cholangiocyte-like subpopulations. e, Expression of Hepatocyte progenitor associated markers in the cholangiocyte dataset.

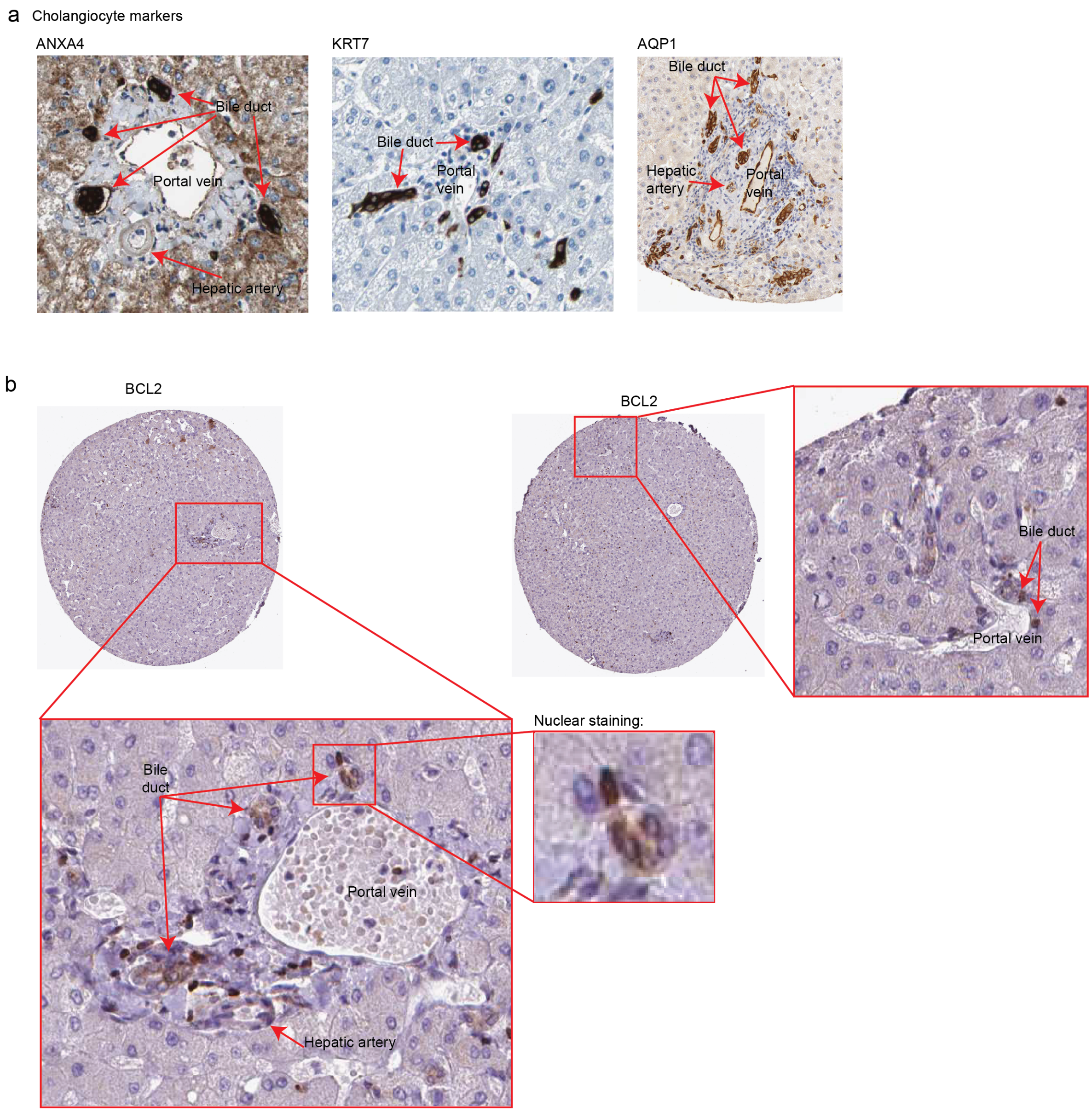


**Extended Data Fig. 14: Bile duct enrichment of cholangiocyte and *BCL2*+ cholangiocyte specific marker proteins via immunohistochemistry from the Human Protein Atlas.** a, immunostaining of cholangiocyte markers (ANXA4, KRT7, AQP1) at the periportal regions indicate expression at bile ducts of non-fibrotic healthy human livers (6). b, BCL2 expression stains the nuclei of cells lining the bile ducts in the healthy human liver through the Human Protein Atlas (6).

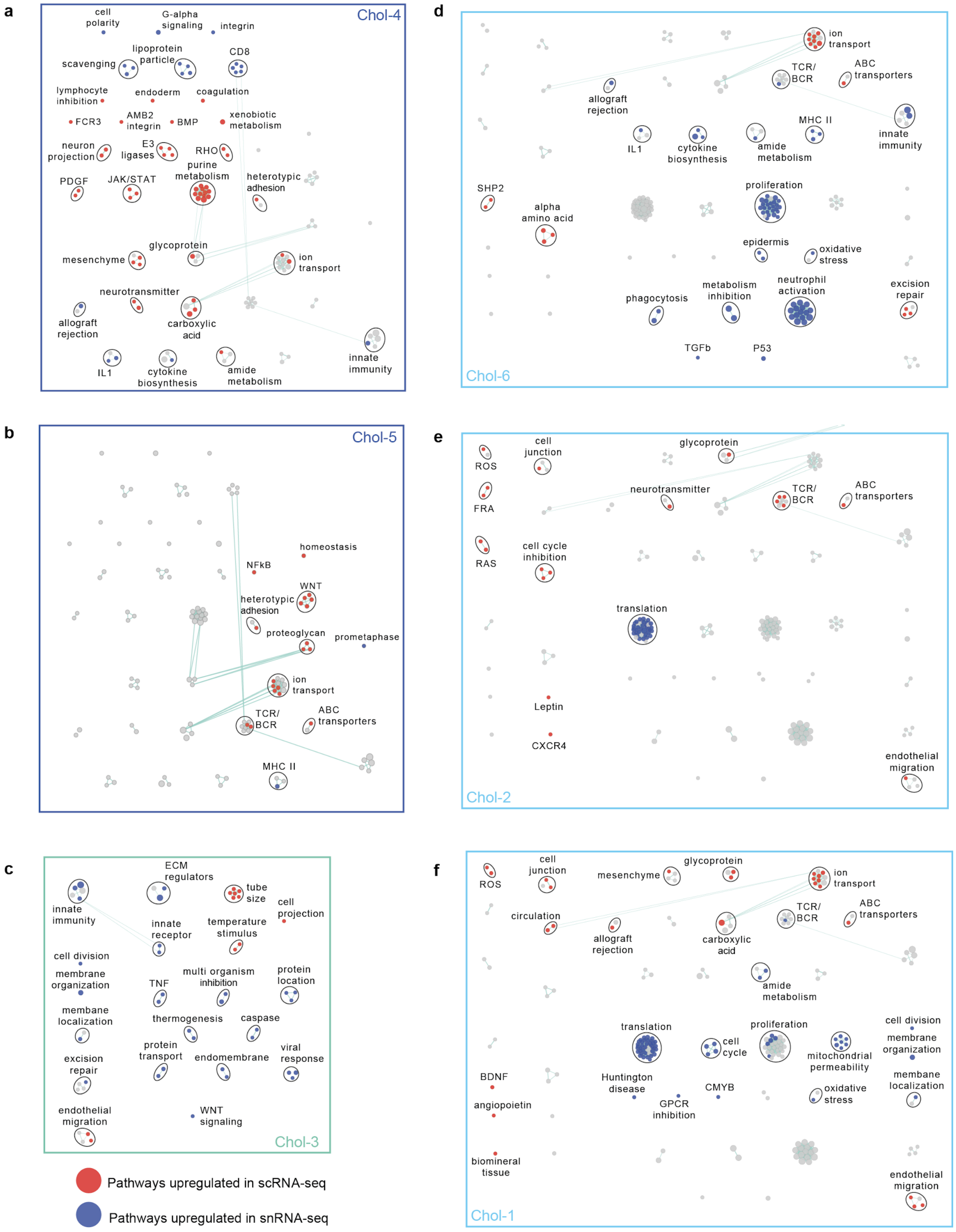
a
Chol-4
cell polarity
G-alpha signaling
integrin
lipoprotein particle
CD8
scavenging
lymphocyte inhibition
endoderm
coagulation
FCR3
AMB2 integrin
BMP
xenobiotic metabolism
neuron projection
E3 ligases
RHO
purine metabolism
PDGF
JAK/STAT
heterotypic adhesion
glycoprotein
mesenchyme
ion transport
neurotransmitter
carboxylic acid
allograft rejection
innate immunity
IL1
cytokine biosynthesis
amide metabolism
b
Chol-5
homeostasis
NFkB
WNT
heterotypic adhesion
proteoglycan
prometaphase
ion transport
TCR/ BCR
ABC transporters
MHC II
c
ECM regulators
tube size
cell projection
innate immunity
innate receptor
temperature stimulus
cell division
membrane organization
TNF
multi organism inhibition
protein location
membane localization
thermogenesis
caspase
excision repair
protein transport
endomembrane
viral response
endothelial migration
WNT signaling
Chol-3
Pathways upregulated in scRNA-seq
Pathways upregulated in snRNA-seq
d
ion transport
TCR/ BCR
ABC transporters
allograft rejection
MHC II
innate immunity
IL1
cytokine biosynthesis
amide metabolism
SHP2
proliferation
alpha amino acid
epidermis
oxidative stress
metabolism inhibition
neutrophil activation
excision repair
phagocytosis
TGFb
P53
Chol-6
e
cell junction
glycoprotein
ROS
FRA
neurotransmitter
TCR/ BCR
ABC transporters
RAS
cell cycle inhibition
translation
Leptin
CXCR4
endothelial migration
Chol-2
f
ROS
cell junction
mesenchyme
glycoprotein
ion transport
circulation
allograft rejection
carboxylic acid
TCR/ BCR
ABC transporters
amide metabolism
translation
proliferation
cell division
cell cycle
mitochondrial permeability
membrane organization
BDNF
Huntington disease
CMYB
oxidative stress
membrane localization
angiopoietin
GPCR inhibition
biomineral tissue
endothelial migration
Chol-1

**Extended Data Fig. 15: Breakdown of pathways enriched in scRNA-seq vs snRNA-seq in the cholangiocyte sub-populations.** a-f, For each population, Fig. 3f is broken down into each cluster. Enriched pathways in scRNA-seq are labeled in red and pathways enriched in snRNA-seq are indicated in blue. Circles (nodes) represent pathways, sized by the number of genes included in that pathway. Related pathways, indicated by light blue lines, are grouped into a theme (black circle) and labeled.

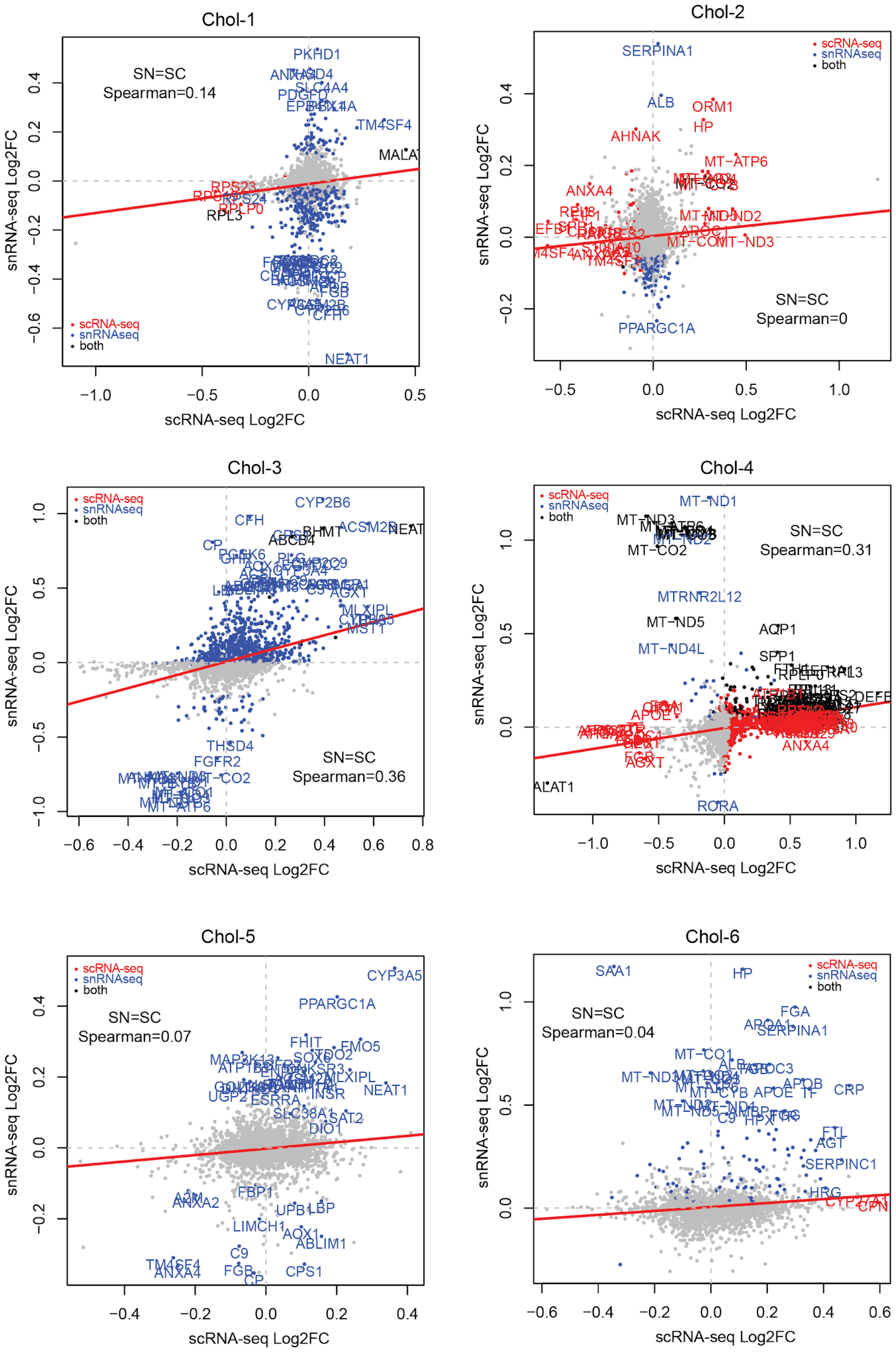


**Extended Data Fig. 16: Differences in upregulated genes between scRNA-seq and snRNA-seq for cholangiocyte subpopulations.** Log2FC of significant genes (5% FDR) within either scRNA-seq (red) or snRNA-seq (blue) or both (black) for each cluster within the cholangiocyte-like sub-clusters, non-significant shown in grey, and Spearman correlation coefficient indicate on the plots.

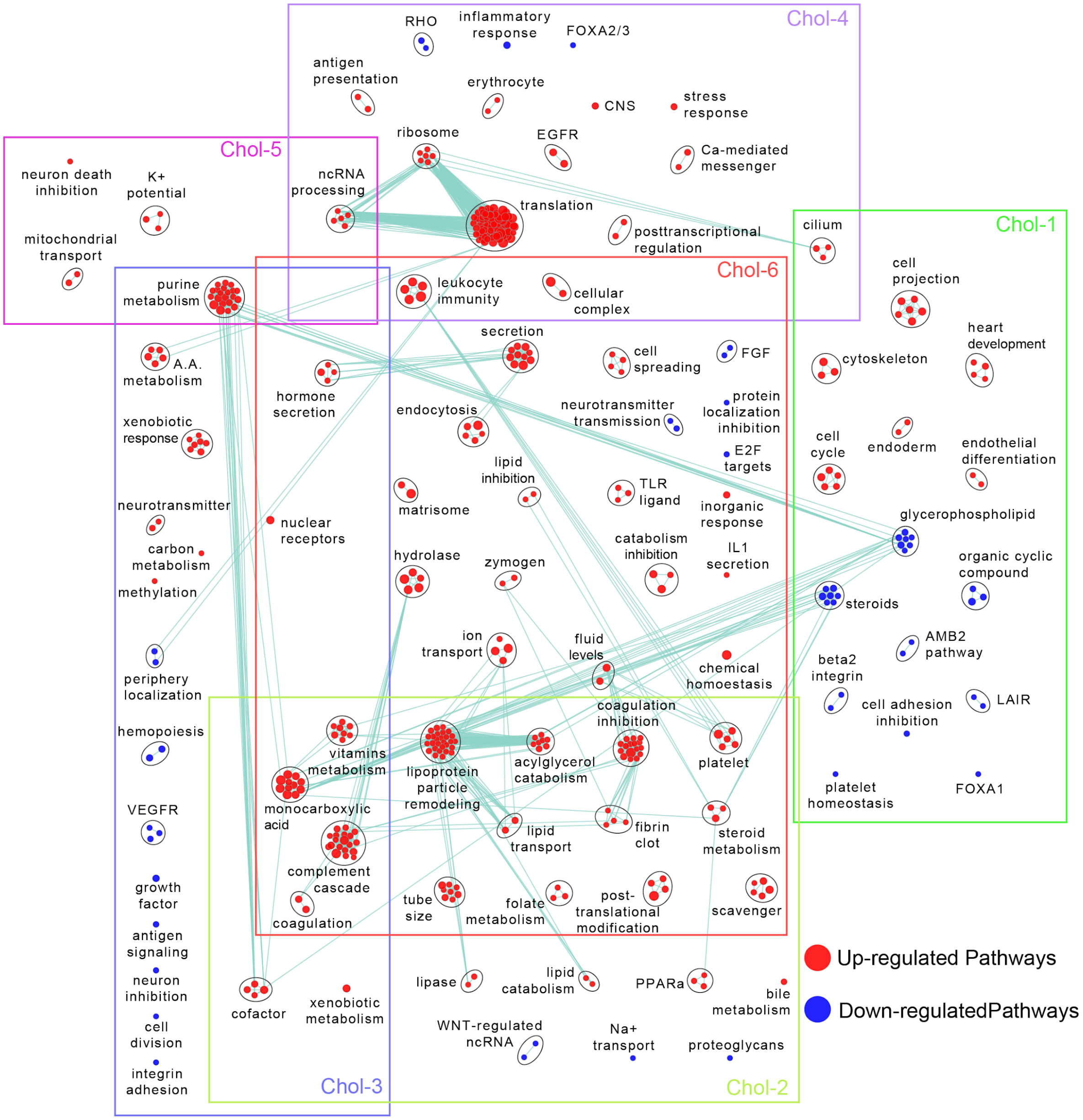


**Extended Data Fig. 17: Pathway enrichment analysis examining active cellular pathways in the cholangiocyte-associated subpopulations.** Up-regulated pathways in each cluster are labeled in red and down-regulated pathways are indicated in blue. Circles (nodes) represent pathways, sized by the number of genes included in that pathway. Related pathways, indicated by light blue lines, are grouped into a theme (black circle) and labeled.

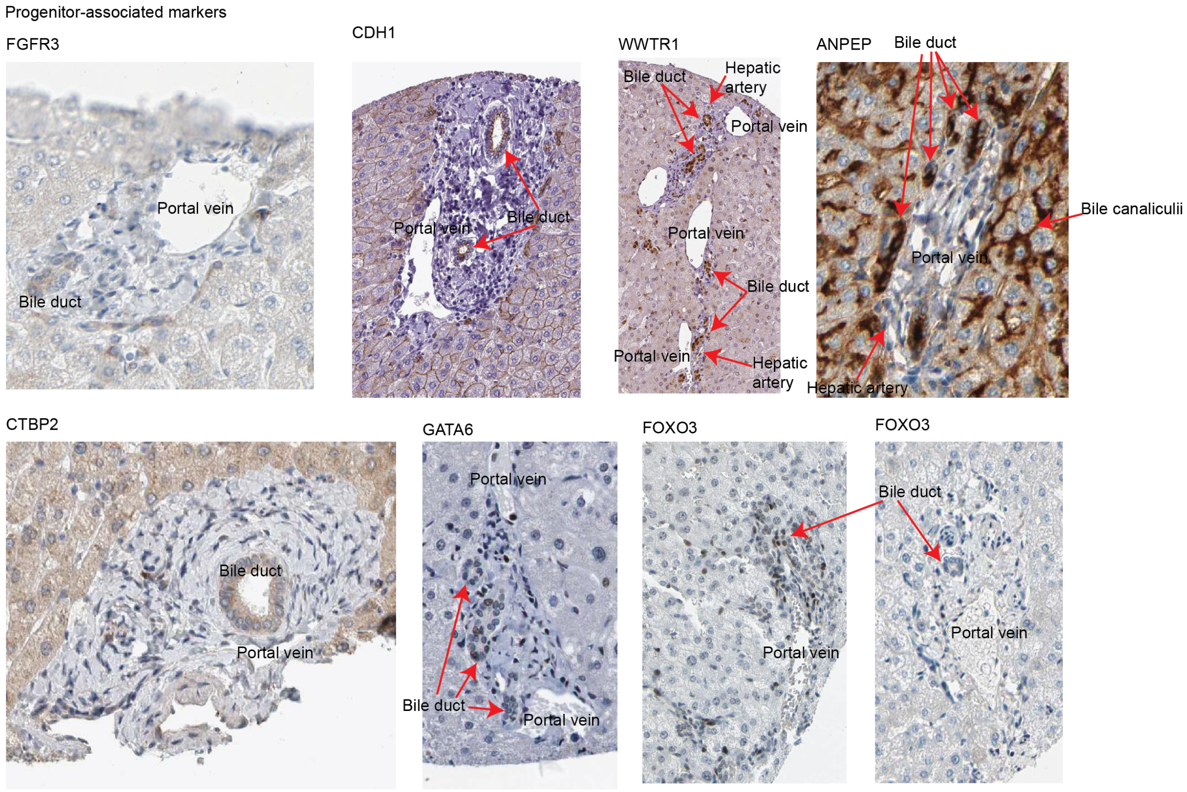


**Extended Data Fig. 18: Progenitor-associated markers immunostaining from the Human Protein Atlas (6).** The images show expressions of each protein at the bile duct. ANPEP stains both the bile ducts and the bile canaliculi. CDH1, an epithelial cell marker, is found on the membranes of hepatocytes and on the luminal side of the bile duct.

a

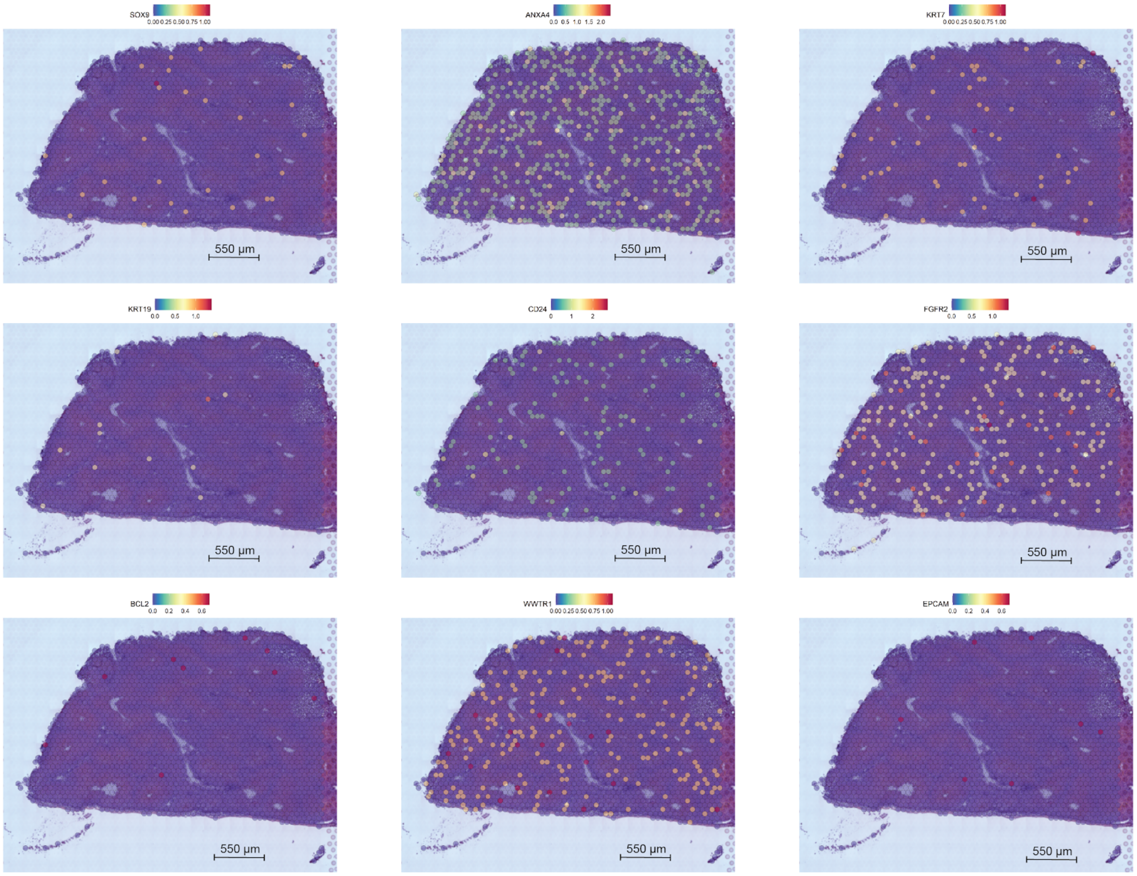
SOX9
ANXA4
KRT7
KRT19
CD24
FGFR2
BCL2
WWTR1
EPCAM
550 µm

b

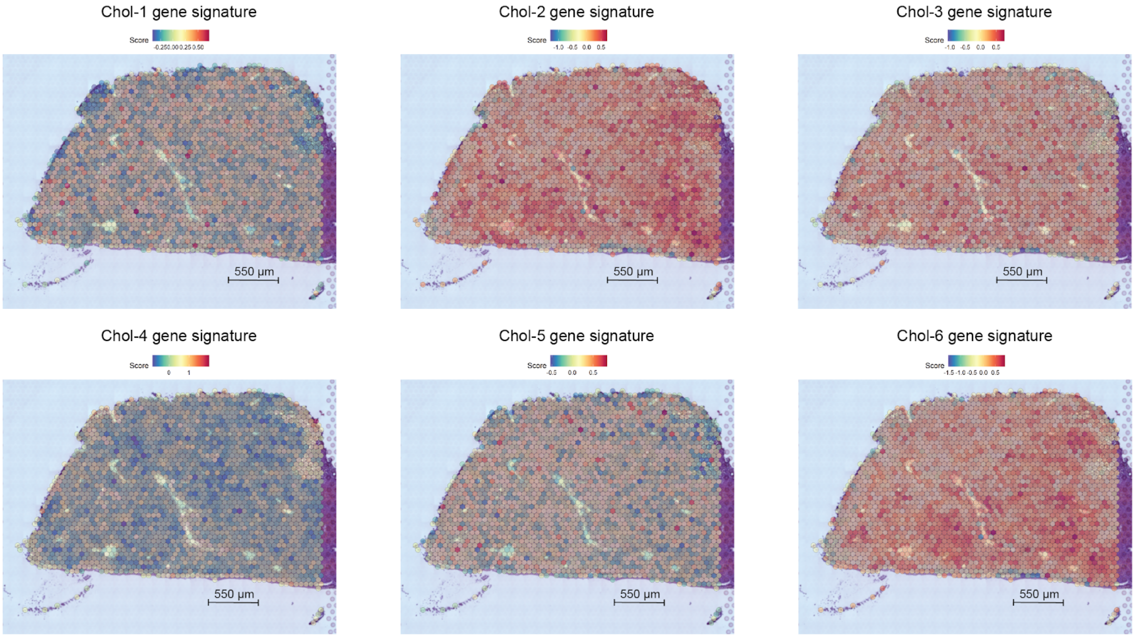
Chol-1 gene signature
Chol-2 gene signature
Chol-3 gene signature
Chol-4 gene signature
Chol-5 gene signature
Chol-6 gene signature
Score
550 µm

**Extended Data Fig. 19: Spatial distribution of cholangiocyte-associated subpopulations across the healthy human liver using spatial transcriptomics (10x genomic Visium platform).** a, Expression of genes upregulated in the cholangiocyte and progenitor clusters in the combined scRNA-seq and snRNA-seq dataset. b, Average expression of the top 30 upregulated genes for each cluster for each cholangiocyte sub-population in the human liver.

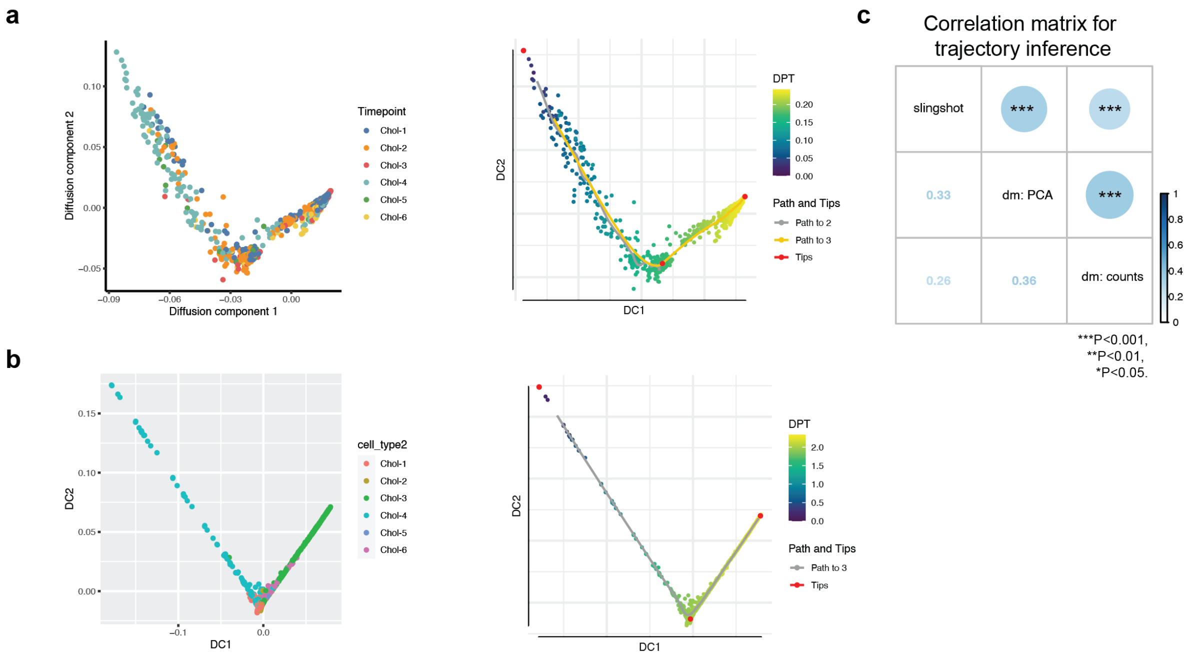


**Extended Data Fig. 20: Cholangiocyte trajectory inference: a, Diffusion maps based on the raw counts matrix.** b, Diffusion maps calculated using the PCA loadings. Cells coloured by cluster (left) and diffusion pseudotime scale (right). Pseudotime scale and predicted paths are indicated on the right. DPT (7) indicates diffusion-pseudotime and is interpreted here as a developmental coordinate. c, Spearman correlation matrix for Slingshot (8) (Fig. 3g) and both diffusion maps trajectory inference indicate significant concordance between all approaches.

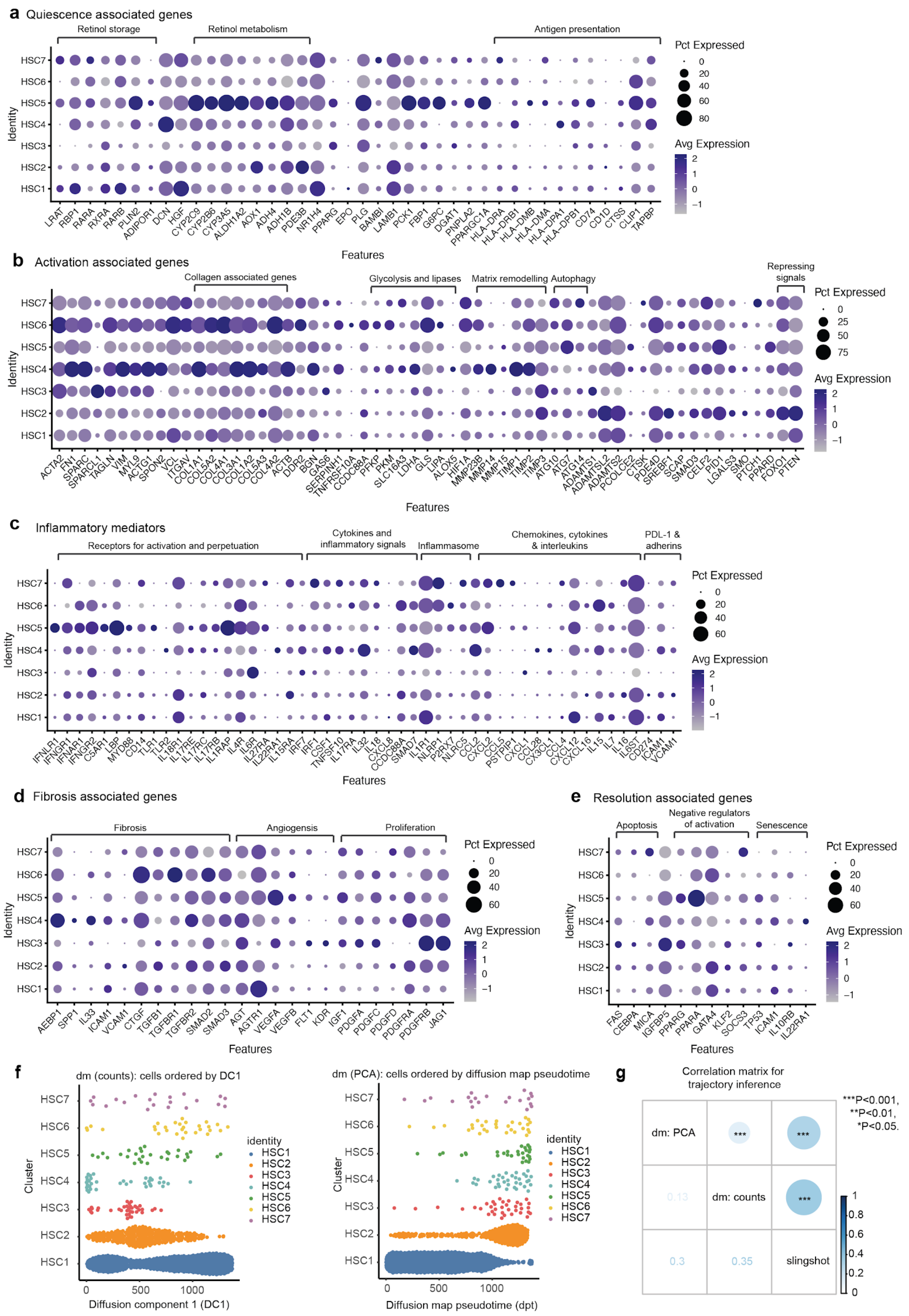
a Quiescence associated genes
Retinol storage
Retinol metabolism
Antigen presentation
Pct Expressed
0
20
40
60
80
Avg Expression
2
1
0
−1
HSC7
HSC6
HSC5
HSC4
HSC3
HSC2
HSC1
Identity
LRAT RBP1 RARA RXRA RARB PLIN2 ADIPOR1 DCN HGF CYP2C9 CYP2B6 CYP3A5 ALDH1A2 AOX1 ADH4 ADH1B PDE3B NR1H4 PPARG EPO PLG BAMBI LAMB1 PCK1 FBP1 G6PC DGAT1 PNPLA2 PPARGC1A HLA-DRA HLA-DRB1 HLA-DMB HLA-DMA HLA-DPA1 HLA-DPB1 CD74 CD1D CTSS CLIP1 TAPBP
Features
b Activation associated genes
Collagen associated genes
Glycolysis and lipases
Matrix remodelling
Autophagy
Repressing signals
Pct Expressed
0
25
50
75
Avg Expression
2
1
0
−1
ACTA2 FN1 SPARC SPARCL1 TAGLN VIM MYL9 ACTG1 SPON2 VCL ITGAV COL1A1 COL5A2 COL4A1 COL3A1 COL1A2 COL5A3 COL4A2 ACTB DDR2 BGN GAS6 SERPINH1 TNFRSF10A CCDC88A PFKP PKM SLC16A3 LDHA GLS LIPA ALOX5 HIF1A MMP23B MMP14 MMP15 TIMP1 TIMP2 TIMP3 ATG10 ATG7 ATG14 ADAMTS1 ADAMTSL2 ADAMTS2 PCOLCE2 CTSK PDE4D SREBF1 SCAP SMAD3 CELF2 PID1 LGALS3 SMO PTCH1 PPARD FOXO1 PTEN
Features
c Inflammatory mediators
Receptors for activation and perpetuation
Cytokines and inflammatory signals
Inflammasome
Chemokines, cytokines & interleukins
PDL-1 & adherins
Pct Expressed
0
20
40
60
Avg Expression
2
1
0
−1
IFNLR1 IFNGR1 IFNAR1 IFNGR2 C5AR1 LBP MYD88 CD14 TLR1 TLR2 IL18R1 IL17RE IL17RC IL17RB IL1RAP IL4R IL6R IL27RA IL22RA1 IL15RA IRF7 IRF1 CSF1 TNFSF10 IL17RA IL32 IL18 CXCL8 CCDC88A SMAD7 IL1R1 NLRP1 P2RX7 NLRC5 CCL2 CXCL2 CCL5 PSTPIP1 CXCL1 CCL28 CX3CL1 CCL4 CXCL12 CXCL16 IL15 IL7 IL16 IL6ST CD274 ICAM1 VCAM1
Features
d Fibrosis associated genes
Fibrosis
Angiogensis
Proliferation
Pct Expressed
0
20
40
60
Avg Expression
2
1
0
−1
AEBP1 SPP1 IL33 ICAM1 VCAM1 CTGF TGFB1 TGFBR1 TGFBR2 SMAD2 SMAD3 AGT AGTR1 VEGFA VEGFB FLT1 KDR IGF1 PDGFA PDGFC PDGFD PDGFRA PDGFRB JAG1
Features
e Resolution associated genes
Apoptosis
Negative regulators of activation
Senescence
Pct Expressed
0
20
40
60
Avg Expression
2
1
0
−1
FAS CEBPA MICA IGFBP5 PPARG PPARA GATA4 KLF2 SOCS3 TP53 ICAM1 IL10RB IL22RA1
Features
f
dm (counts): cells ordered by DC1
Cluster
0 500 1000
Diffusion component 1 (DC1)
identity
HSC1
HSC2
HSC3
HSC4
HSC5
HSC6
HSC7
dm (PCA): cells ordered by diffusion map pseudotime
0 500 1000
Diffusion map pseudotime (dpt)
g
Correlation matrix for trajectory inference
dm: PCA
***
***
0.13
dm: counts
***
0.3
0.35
slingshot
***P<0.001,
**P<0.01,
*P<0.05.
1
0.8
0.6
0.4
0.2
0

**Extended Data Fig. 21: Hepatic stellate cell markers and trajectory inference.** DotPlots indicate the relative average expression of HSC markers of quiescence (a), activation (b), inflammation (c), fibrosis (d), and clearance (e) from the literature across the stellate cell subpopulations in the combined scRNA-seq and snRNA-seq dataset. f, Diffusion maps based on the raw counts matrix (left) and Diffusion maps calculated using the PCA loadings (right). Cells coloured by cluster (left) and ordered along the X axis as a measure of diffusion-pseudotime and is interpreted here as a developmental coordinate. g, Spearman correlation matrix for Slingshot (8) (Fig. 4g) and both diffusion maps trajectory inference indicate significant concordance between all approaches.

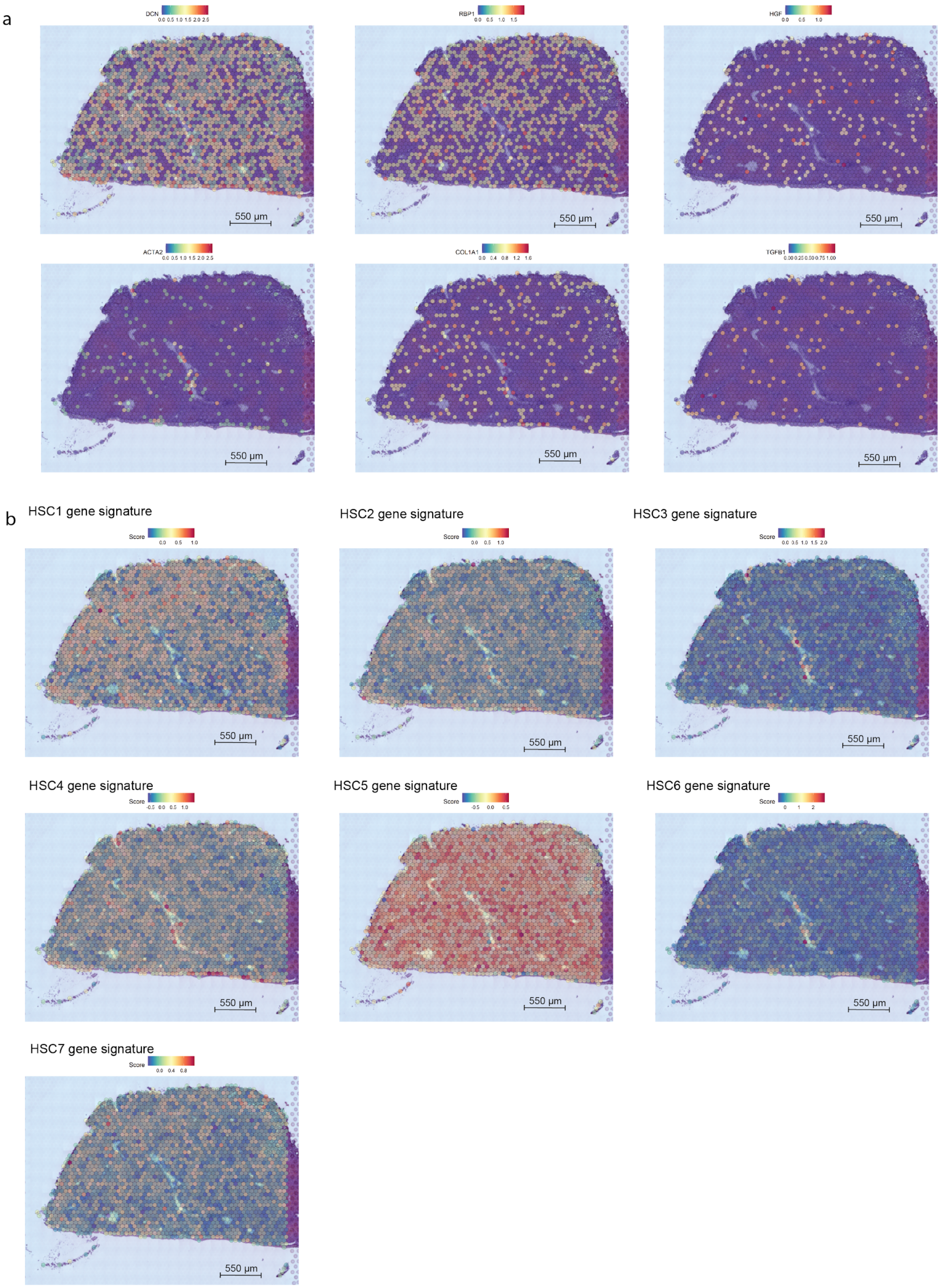
a
DCN
RBP1
HGF
ACTA2
COL1A1
TGFB1
550 µm
b
HSC1 gene signature
HSC2 gene signature
HSC3 gene signature
HSC4 gene signature
HSC5 gene signature
HSC6 gene signature
HSC7 gene signature
Score

**Extended Data Fig. 22: Hepatic stellate cell genes and gene signatures in the healthy human liver using spatial transcriptomics (10x genomic Visium platform).** a, Quiescent HSC (*DCN, RBP1, HGF*) and activated HSC (*ACTA2, COL1A1, TGFB*) markers across healthy human liver tissue. b, For each HSC cluster, the average expression of the top 30 genes (gene signature) overlaid on a slice of healthy human liver tissue.

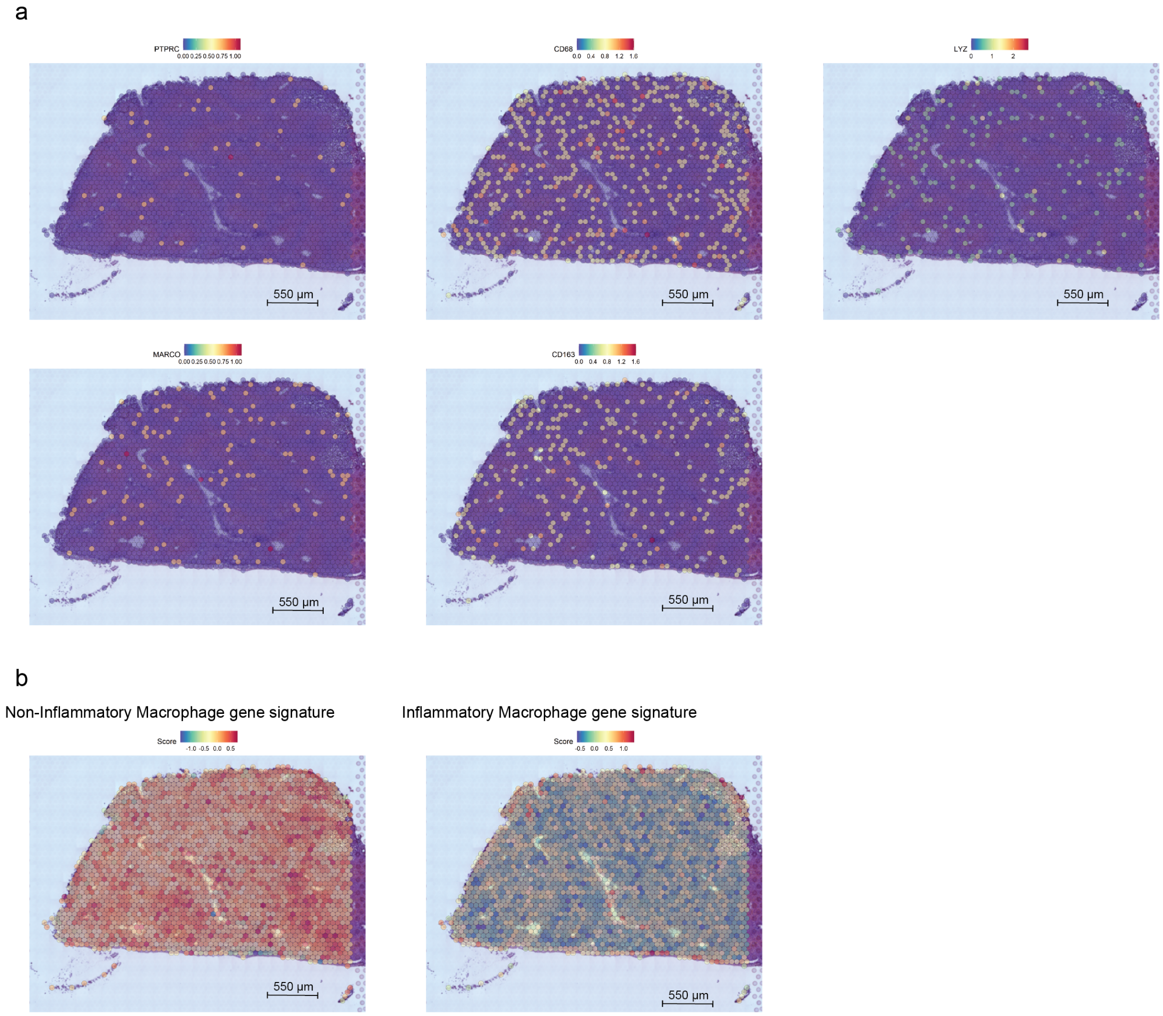


**Extended Data Fig. 23: Verification of macrophage clusters in the healthy human liver using spatial transcriptomics (10x genomic Visium platform).** a, Expression of immune-cell marker *PTPRC* (CD45) and hepatic macrophage markers (*CD86, LYZ, MARCO, CD163*). b, Inflammatory and non-inflammatory macrophage gene signatures (Top 30 genes per cluster) in the spatial RNA data.

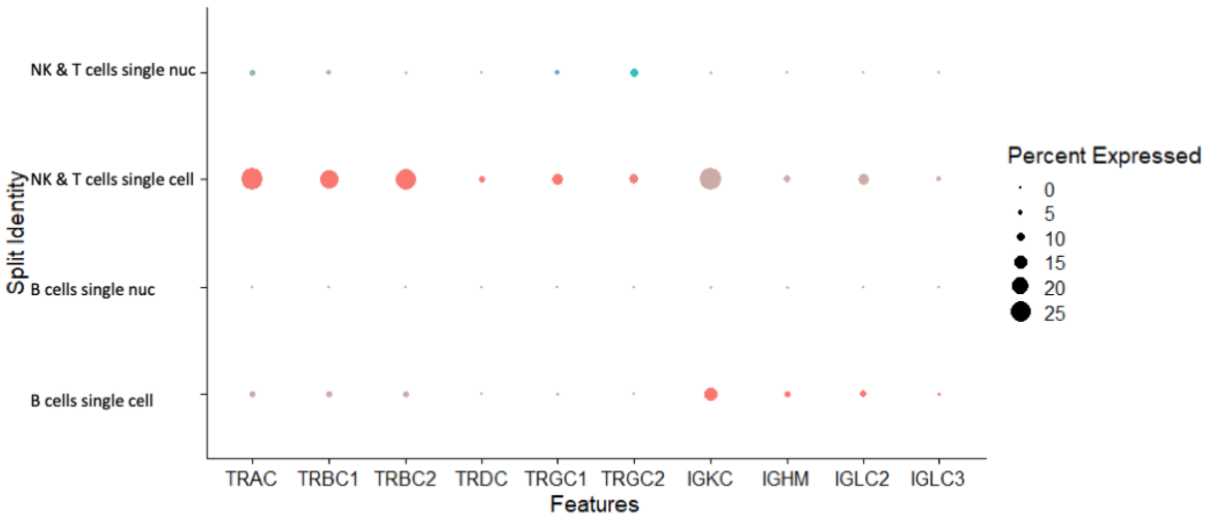


**Extended Data Fig. 24: Expression of TCR and BCR components in scRNA-seq and snRNA-seq.**

**Supplementary Table 1: Core cell annotation markers-** available as a dropbox link due to large file size
https://www.dropbox.com/s/0sf67ni18cuqujy/Supplementary%20Table%201.xlsx?dl=0

| | **scRNA-seq (n=29,432)** | **snRNA-seq (n=43,863)** |
|---|---|---|
| **13 MT RNA** | 21±12% | 3±5% |
| **89 Ribosomal RNA/Proteins** | 12±9% | 0.4±0.4% |
| **gene/umi** | 0.42±0.13 | 0.51±0.17 |
| **gene/umi - exclude MT & Ribo** | 0.55±0.14 | 0.52±0.18 |

**Supplementary Table 2: RNA composition of scRNA-seq and snRNA-seq - mean±SD**
Single-cell is less sensitive than single nuc because of the high proportion of mitochondrial and ribosomal-related RNA.

| **Donor** | **scRNA-seq** | **snRNA-seq: TST** | **snRNA-seq: CST** | **snRNA-seq: NST** |
|---|---|---|---|---|
| **C41** | 740±36 umis<br>346±7 genes | 2348±30 umis<br>1407±10 genes | 2389±62 umis<br>1435±15 genes | 3953±97 umis<br>2191±19 genes |
| **C58** | 2272.5±88 umis<br>832±12 genes | 565±13 umis<br>389±5 genes | NA | NA |
| **C70** | 987±163 umis<br>342±12 genes | 7838±77 umis<br>3037±10 genes | NA | NA |
| **C72** | 976±19 umis<br>505±4 genes | 8525±58 umis<br>3109±9 genes | NA | NA |

**Supplementary Table 3: Median UMI counts and number of detected genes per cell / nucleus across all samples along with standard errors across cells within each sample.**

| Cell Type | scRNA-seq | snRNA-seq | snRNA-seq: CST | snRNA-seq:NST | snRNA-seq: TST |
|---|---|---|---|---|---|
| **Hepatocyte** | 71.7±5.7% | 67.9±6.7% | 47.5% | 52.4% | 76.9±5.6% |
| **LSEC** | 8.9±2.2% | 12.8±3.4% | 25.1% | 19.7% | 8.0±2.3% |
| **Macrophage** | 6.13±2.0% | 5.8±1.4% | 9.1% | 8.2% | 4.3±1.7% |
| **T and NK-like** | 4.5±1.7% | 3.1±1.1% | 7.2% | 1.4% | 2.5±1.2% |
| **HSC** | 3.7±1.0% | 6.0±1.3% | 6.5% | 12.1% | 4.3±0.5% |
| **B cells** | 2.7±1.6% | 1.1±0.2% | 1.3% | 0.5% | 1.2±0.2% |
| **Cholangiocyte** | 2.4±0.9% | 3.4±0.6% | 3.3% | 5.7% | 2.8±0.7% |
| **Erythroid-like** | 0.01±0.005 % | 0.01±0.006% | 0 | 0 | 0.01±0.01% |

**Supplementary Table 4: Cell type compositions for scRNA-seq and snRNA-seq.**
Mean cell type frequency and standard errors across the four donor samples. Only one sample was sequenced with CST and NST thus standard errors could not be calculated.

| Cluster | scRNA-seq | snRNA-seq | % of total cells in scRNA-seq map | % of total cells in snRNA-seq map |
|---|---|---|---|---|
| **CV1** | 2575 | 7608 | 8.749 | 17.345 |
| **IZ1** | 4505 | 3637 | 15.306 | 8.292 |
| **PP1** | 4756 | 12475 | 16.159 | 28.441 |
| **PP2** | 4685 | 4166 | 15.918 | 9.498 |
| **Unidentified** | 576 | 1309 | 1.957 | 2.985 |
| **IZ2** | 2693 | 1274 | 9.150 | 2.905 |
| **All Hepatocytes** | 19790 | 30469 | 67.240 | 69.4648 |

**Supplementary Table 5: Frequency of hepatocyte clusters in scRNA-seq and snRNA-seq**

| Cluster | scRNA-seq | snRNA-seq | % of total cells in scRNA-seq map | % of total cells in snRNA-seq map |
|---|---|---|---|---|
| **Chol-1** | 134 | 621 | 0.455 | 1.416 |
| **Chol-2** | 146 | 169 | 0.496 | 0.385 |
| **Chol-3** | 32 | 234 | 0.109 | 0.533 |
| **Chol-4** | 90 | 145 | 0.306 | 0.331 |
| **Chol-5** | 33 | 177 | 0.112 | 0.404 |
| **Chol-6** | 13 | 67 | 0.044 | 0.153 |
| **All Cholangiocytes** | 448 | 1413 | 1.522 | 3.22 |

**Supplementary Table 6: Representation of cholangiocyte clusters in scRNA-seq and snRNA-seq**

| Cluster | scRNA-seq | snRNA-seq | % of total cells in scRNA-seq map | % of total cells in snRNA-seq map |
|---|---|---|---|---|
| **HSC1** | 202 | 703 | 0.686 | 1.603 |
| **HSC2** | 65 | 254 | 0.221 | 0.579 |
| **HSC3** | 8 | 28 | 0.027 | 0.064 |
| **HSC4** | 17 | 18 | 0.058 | 0.041 |
| **HSC5** | 4 | 25 | 0.0136 | 0.0570 |
| **HSC6** | 4 | 25 | 0.0136 | 0.0570 |
| **HSC7** | 5 | 16 | 0.0170 | 0.0365 |
| **All HSC** | 305 | 1069 | 1.036 | 2.437 |

**Supplementary Table 7: HSC recovery in scRNA-seq vs. snRNA-seq.**

| Cluster | scRNA-seq | snRNA-seq | % of total cells in scRNA-seq map | % of total cells in snRNA-seq map |
|---|---|---|---|---|
| **Central Venous LSEC** | 1552 | 3299 | 5.273 | 7.521 |
| **Periportal LSEC** | 340 | 579 | 1.155 | 1.320 |
| **Portal Endothelial** | 186 | 333 | 0.632 | 0.759 |
| **All Endothelial cells** | 2078 | 4211 | 7.060 | 9.600 |

**Supplementary Table 8: Capture of LSEC subtypes by each technology.**

| Cluster | scRNA-seq | snRNA-seq | % of total cells in scRNA-seq map | % of total cells in snRNA-seq map |
|---|---|---|---|---|
| **Inflammatory** | 1279 | 828 | 4.346 | 1.888 |
| **Non-Inflammatory** | 909 | 961 | 3.088 | 2.191 |
| **All Macrophages** | 2188 | 1789 | 7.434 | 4.079 |

**Supplementary Table 9: Frequency of macrophage subpopulations across scRNA-seq and snRNA-seq.**

| Cluster | scRNA-seq | snRNA-seq | % of total cells in scRNA-seq map | % of total cells in snRNA-seq map |
|---|---|---|---|---|
| **αβ T cells** | 936 | 458 | 3.180 | 1.044 |
| **NK cells** | 254 | 162 | 0.863 | 0.369 |
| **ɣδ T cells** | 209 | 93 | 0.710 | 0.212 |
| **Mature B cells** | 81 | 44 | 0.275 | 0.100 |
| **Plasma B cells** | 62 | 47 | 0.211 | 0.107 |
| **All Lymphocytes** | 1542 | 804 | 5.239 | 1.833 |
| **All B cells** | 143 | 91 | 0.486 | 0.207 |

**Supplementary Table 10: Recovery of lymphocyte populations by scRNA-seq and snRNA-seq**

**Supplementary Table 11: Cell Ranger summary - please see attached.**

**Supplementary Table 12: SoupX genes - please see attached.**

### EXTENDED EXPERIMENTAL PROCEDURES

#### Preparation of fresh tissue homogenates for single cell RNA sequencing

Human liver tissue from the caudate lobe was obtained from livers procured from neurologically deceased donors deemed acceptable for liver transplantation. Samples were collected with appropriate institutional ethics approval from the University Health Network (REB# 14-7425-AE). A fragment of the liver tissue was preserved for snRNA-seq by snap freezing in liquid nitrogen. Single cell suspensions of fresh human liver were generated as previously described (9). Briefly, during organ retrieval, donor grafts were perfused in situ with cold (HTK) solution (Methapharm) to flush circulating cells. The remaining tissue resident parenchymal and non-parenchymal cells were then used to prepare a single-cell suspension for scRNA-seq analysis. At our institute, the caudate lobe (segment 1) of the liver is often removed in preparation of the organ for implantation. The removed caudate lobe was then dissociated using the two step collagenase perfusion protocol [https://doi.org/10.17504/protocols.io.m9sc96e].

**Isolation of nuclei from snap-frozen human liver tissue**

Single nucleus extraction was performed as previously described (10). Briefly, nuclei were isolated from fresh frozen tissue using the salt-tris (ST) based buffer [73 mM NaCl (Sigma-Aldrich, St Louis, MO, USA), 5 mM Tris-HCl pH 7.5 (Invitrogen, Grand Island, NY, USA), 0.5 mM CaCl2 (Sigma-Aldrich, St Louis, MO, USA), and 10.5 mM MgCl2 (Thermo Fisher Scientific, Vilnius, LT), 0.01% BSA (New England Biolabs, Ipswich, MA, USA) ] containing 0.03% Tween-20 (BioShop, Burlington, ON, CA) (TST), 0.5% CHAPS (CST) (Sigma-Aldrich, St Louis, MO, USA), or 0.2% NonidetTM P40 Substitute (NST) (Roche, Mannheim, Germany). Samples (< 3 mm pieces) were lysed in 1 mL of ice-cold TST/CST/NST using spring scissors (14 mm cutting edge, 0.275 mm tip diameter, 12 cm length) (Fine Science Tools) for 10 minutes. The homogenate was filtered through a 40 µm cell strainer (Falcon, Corning, NY, USA) and washed with 1 mL of TST/CST/NST followed by 3 mL of ice-cold ST buffer. The nuclei suspension was centrifuged at 500 *x g* at 4ºC for 5 min with no brake. The pellet was resuspended in 0.01% BSA ST buffer and filtered through a 40 µm cell strainer. The number and integrity of the nuclei and the purity of the preparation were determined using bright-field microscopy and fluorescence microscopy for RNA visualization with SYBR Green II RNA gel stain (Invitrogen, Grand Island, NY, USA).

**10x sample processing, cDNA library preparation and sequencing**

Samples were prepared as outlined by the 10x Genomics Single Cell 3′ v2 and 3' v3 Reagent Kit user guides and as described previously (9). Briefly, following cell counting (using Trypan blue exclusion or SYBR Green II RNA gel stain), we targeted capture of 6000 cells or 6000 nuclei and loaded onto the 10x Genomics single-cell-A or B chip. cDNA libraries were prepared as per the Single Cell 3′ Reagent Kits v2 or v3 user guide. Samples were sequenced on a HiSeq 2500 or NovaSeq 6000. Sequencing QC summaries for each liver profiled are found in Supplementary Table 11. Due to timing of sample acquisition, the snRNA-seq samples were processed with v3 chemistry and scRNA-seq samples were processed with v2 chemistry. V3 chemistry differs from v2 chemistry in two ways (11). V3 is biased towards higher mitochondrial reads which suggests our estimate of the mitochondrial fraction in snRNA-seq is an overestimate relative to the fraction in scRNA-seq. V3 captures more UMIs relative to the total reads sequenced than v2. As all of the statistics in this manuscript are always relative to the total number of UMIs per cell/nucleus this difference does not impact our results.

**DNA sequencing data processing**

Single cell data was processed using 10x Cell Ranger software version 3.01, mapping reads to the GRCh38 human genome. Single nucleus data was processed using Cell Ranger version 3, and reads were mapped to a modified transcriptome based on GRCh38 which included intronic regions to ensure quantification of reads derived from immature, unspliced mRNA present in the nucleus. Droplets containing fewer than 10 total unique molecular identifiers (UMIs) were excluded, then cells and nuclei were called from the raw count matrices using EmptyDrops with 5% FDR (12). We also excluded all genes that were detected in fewer than 10 cells or nuclei, cells or nuclei with fewer than 100 detected genes, and cells and nuclei with greater than 50% of

UMIs mapped to the mitochondrial genome. Each sample was size-factor normalized to 10,000 UMI per cell/nucleus, then log2 transformed and scaled using Seurat (version 3.1.3), the top 2000 most highly variable features in each sample were identified using Seurat. Ambient RNA was estimated using SoupX (13), version 1.2.2, using the supervised approach employing known cell-type specific markers (Supplementary Table 12). However, systematic technical differences between single-cell and single-nucleus sequencing would also appear in the ambient RNA, and SoupX can have variable performance on different samples, thus we did not remove the estimated ambient signal from the samples. Removing ambient RNA does not change our main findings.

**Data integration and clustering**

All samples were merged and subset to the set of genes detected in at least 10 cells or nuclei across all samples. Consensus highly variable genes were identified as those that were found to be highly variable in at least two of the samples. Genes located on the mitochondrial genome were excluded from the list of highly variable genes. This left 2,143 highly variable genes.

The top 15 principal components were identified from the merged individually scaled datasets. The data was integrated using default parameters of Harmony (14), then clustered using Seurat's SNN-Louvain clustering algorithm (15). The data was clustered using 30 sets of parameters, and the most consistent clusterings were identified using apcluster (16) on the cluster-cluster distance matrix calculated using the Variation of Information criterion (17).

Clusters were first automatically annotated using a hypergeometric test, for the enrichment of known markers from our previous human liver map (9), among the top 200 markers for each cluster as identified with Seurat's implementation of the Wilcoxon-rank-sum test. These annotations were further refined through manual curation and correlation with mouse and human hepatic lobule zonation patterns (5,18).

**Sub-clustering and differential gene expression**

Each major population was sub-clustered, scaled individually, merged and Harmony integrated. After clustering, differentially expressed genes were calculated and contaminating clusters were identified and removed according to known markers (Hepatocytes: *ALB*, *HAL, CYP3A4*; LSECs: *CALCRL, VWF, FCGR2B*; Cholangiocytes: *SOX9, KRT7, ANXA4*; Stellate cells: *ACTA2, DCN, RBP1*). The remaining cells were then re-analyzed and annotated based on their expression of cell-type associated markers from the literature. Marker genes for the populations identified in the combined scNRA-seq and snRNA-seq dataset were calculated by performing a pairwise t-test between the genes in each pair of clusters within each sample and subsequently combining into a single p-value using the block argument of scran's findMarkers (19). Seurat's Wilcoxon-rank-sum test was used to calculate differentially upregulated genes for each cluster that are specific to each protocol.

**Gene-type and gene length biases between scRNA-seq and snRNA-seq data**

3,804 housekeeping genes were obtained from the literature (20). Long non-coding RNAs and protein coding gene lists were obtained from Ensembl. Nuclear-encoded mitochondrial proteins were obtained from MitoCarta3.0 (21). Ribosomal genes were obtained from the Ribosomal Protein Gene database (22). Transcript GC content, miRNA binding sites, transcript length, UTR lengths and intron length were obtained from Ensembl Biomart. Log Fold changes of mean expression across all cell-types in scRNA-seq and snRNA-seq were calculated across all samples.

**Pathway enrichment and correlation analysis**

Pathway enrichment analysis was performed as previously described (9) with the addition of a dissociation signature to the pathway gene set database (23). The Human_GOBP_AllPathways_no_GO_iea_November_01_2020_symbol.gmt pathway gene set database from [http://baderlab.org/GeneSets] was used to identify enriched cellular pathways in GSVA and GSEA analysis. Highly related pathways were grouped into a theme and labeled by AutoAnnotate (Version 1.2) in Cytoscape (Version 3.8.2). GSEA and GSVA results were visualized using the Enrichment Map app (Version 3.3) in Cytoscape (Version 3.8.2) (24). Spearman's rank correlation between each hepatocyte cluster and Human/mouse laser capture microdissection bulk RNA-seq was performed as previously published (5,18).

**Trajectory inference analysis**

Slingshot (Version 1.8.0) was employed to infer the pseudotime based on the Harmony embedding matrix of cells. Lineages were calculated using the Slingshot UMAP embedding protocol (8). Diffusion maps (7) (Version 3.1.1) were computed with both, the raw counts matrix as well as the PCA loadings. Spearman's rank correlation coefficient was calculated on each pair of outputs of these analyses and plotted using corrplot (Version 0.84).

**Validation of zonated gene signatures using spatial transcriptomics**

Healthy human liver tissue was embedded in OCT, frozen and cryosectioned with 16um thickness at −10C (cryostar NX70 HOMP). Sections were placed on a chilled Visium Tissue Optimization Slide (10x Genomics) and processed following the Visium Spatial Gene Expression User Guide. Briefly, tissue was permeabilized for 12 minutes, based on an initial optimizations trial and libraries were prepared according to the Visium Spatial Gene Expression User Guide. Samples were sequenced on a NovaSeq 6000.

**Visium spatial transcriptomics data processing**

The Visium spatial transcriptomic data was sequenced to a depth of 167,400,637 reads, a saturation of 77%. These reads were mapped to the GRCh38 human genome and expression was quantified with the spaceranger-1.1.0. Further processing and visualization was performed with Seurat (version 3.2.1). The tissue slice covered 2,265 unique spots with a median of 1,884 genes detected and 6,010 transcripts per spot. This data was normalized to 20,000 UMI per spot and log2 transformed, then each gene was scaled across all spots.

Genes were selected from the significantly differentially expressed genes (FDR < 5%), ranked by the fold change in expression in each cluster or subcluster relative to all other clusters. The top 10 differentially expressed genes in the snRNA-seq, scRNA-seq, and the combined data used for the gene signatures of each cluster. Genes repeated across both or all data types were weighted accordingly, then the average scaled expression across all genes was calculated as the signature score for each spot.

**Validation of zonated protein expression *via* the Human Protein Atlas.** Immunostaining images were obtained from the Human Protein Atlas (https://www.proteinatlas.org) (6). Lobule annotation was confirmed by a liver pathologist (C. Thoeni).

https://kb.10xgenomics.com/hc/en-us/articles/360026501692-Do-we-see-a-difference-in-expression-profile-of-3-Single-Cell-v3-chemistry-as-compared-to-v2-chemistry

12. Lun ATL, Riesenfeld S, Andrews T, Dao TP, Gomes T, participants in the 1st Human Cell Atlas Jamboree, et al. EmptyDrops: distinguishing cells from empty droplets in droplet-based single-cell RNA sequencing data. Genome Biol. 2019;20:63.
13. Young MD, Behjati S. SoupX removes ambient RNA contamination from droplet-based single-cell RNA sequencing data. Gigascience. 2020;9.
14. Korsunsky I, Millard N, Fan J, Slowikowski K, Zhang F, Wei K, et al. Fast, sensitive and accurate integration of single-cell data with Harmony. Nat. Methods. 2019;16:1289–1296.
15. Satija R, Farrell JA, Gennert D, Schier AF, Regev A. Spatial reconstruction of single-cell gene expression data. Nat. Biotechnol. 2015;33:495–502.
16. Frey BJ, Dueck D. Clustering by passing messages between data points. Science. 2007;315:972–976.
17. Meilă M. Comparing Clusterings by the Variation of Information. In: Schölkopf B, Warmuth MK, editors. Learning Theory and Kernel Machines. Berlin, Heidelberg: Springer Berlin Heidelberg; 2003. p. 173–187.
18. Halpern KB, Shenhav R, Matcovitch-Natan O, Toth B, Lemze D, Golan M, et al. Single-cell spatial reconstruction reveals global division of labour in the mammalian liver. Nature. 2017;542:352–356.
19. Lun ATL, Bach K, Marioni JC. Pooling across cells to normalize single-cell RNA sequencing data with many zero counts. Genome Biol. 2016;17:75.
20. Eisenberg E, Levanon EY. Human housekeeping genes, revisited. Trends Genet. 2013;29:569–574.
21. Rath S, Sharma R, Gupta R, Ast T, Chan C, Durham TJ, et al. MitoCarta3.0: an updated mitochondrial proteome now with sub-organelle localization and pathway annotations. Nucleic Acids Res. 2021;49:D1541–D1547.
22. Nakao A, Yoshihama M, Kenmochi N. RPG: the Ribosomal Protein Gene database. Nucleic Acids Res. 2004;32:D168-70.
23. van den Brink SC, Sage F, Vértesy Á, Spanjaard B, Peterson-Maduro J, Baron CS, et al. Single-cell sequencing reveals dissociation-induced gene expression in tissue subpopulations. Nat. Methods. 2017;14:935–936.
24. Merico D, Isserlin R, Stueker O, Emili A, Bader GD. Enrichment map: a network-based method for gene-set enrichment visualization and interpretation. PLoS ONE. 2010;5:e13984.
